# Distant homologies and domain conservation of the Hereditary Spastic Paraplegia protein SPG11/ALS5/spatacsin

**DOI:** 10.1101/2020.03.08.982389

**Authors:** Alexander L Patto, Cahir J O’Kane

## Abstract

Loss-of-function mutations in SPG11 protein (spatacsin) are a common cause of autosomal recessive hereditary spastic paraplegia with thin corpus callosum. To identify regions of the protein that may have functions that are disrupted in disease, we carried out bioinformatic analyses of its conserved regions. An N-terminal region of around 650 amino-acid residues, present in SPG11 across a wide range of metazoan animals, was missing from many insect lineages. Evolutionary loss of this domain correlated with loss of its binding partner, the AP-5 adaptor complex, suggesting that its main function is interaction with AP-5 in intracellular trafficking, and that the remainder of SPG11 carries out AP-5-independent functions. At the C-terminus of SPG11, a spatacsin_C domain showed sequence similarity and predicted structural homology to the Vps16_C domain of the HOPS complex protein Vps16. It localized to acidic compartments, consistent with a role in endolysosomal or autolysosomal transport, like Vps16. Mass spectrometry analysis of binding partners of this domain identified membrane trafficking proteins, some SM proteins, and several aminoacyl-tRNA synthetases. Since mutations affecting SPG11 or aminoacyl-tRNA synthetases can both cause Charcot-Marie-Tooth neuropathy (CMT) type 2, we suggest autolysosomal trafficking as a target process in CMT type 2.

## Introduction

Loss-of-function mutations in *SPG11* commonly result in autosomal recessive Hereditary Spastic Paraplegia (ARHSP) with thinning of the corpus callosum (TCC) (Stevanin *et al*. 2007). *SPG11* is the most common recessive HSP, with HSP-causing mutations identified throughout the coding region. *SPG11* mutations can also cause other degenerative disorders including juvenile ALS (Orlacchio *et al*. 2010; Daoud *et al*. 2012), juvenile Parkinson’s disease and dopaminergic neuron loss (Guidubaldi *et al*. 2011; Anheim *et al*. 2009), Kjellin’s syndrome, or the peripheral neuropathy Charcot-Marie-Tooth disease (Fridman and Murphy 2014; Montecchiani *et al*. 2016).

Loss of *SPG11* in mammals leads to axonal pathology, including axon outgrowth defects in mice (Pérez-Brangulí *et al*. 2014). Young *Spg11* knockout mice have severe neuronal loss in the cerebellum, motor cortex and Purkinje cells (Varga *et al*. 2015). Externally, mice develop a progressive gait disorder, developing ataxia after 12 months (Varga *et al*. 2015), similar to symptoms of *SPG11* patients (Zhao *et al*. 2013). In zebrafish, severe knockdown of *Spg11* or *Spg15* led to similar defects in spinal motor neuron axon outgrowth and locomotion difficulties (Martin *et al*. 2012). *Spg11* phenotypes in animal models and humans are broadly consistent, suggesting important roles for SPG11 in neuronal development and function.

SPG11 localizes to Lamp1-positive compartments in human fibroblasts (Hirst *et al*. 2013), which comprise lysosomes (Rohrer *et al*. 1996), autolysosomes (Yu *et al*. 2010) and late endosomes (Szymanski *et al*. 2011). SPG11 and its binding partner SPG15 show structural homology with clathrin heavy chain and COP1 subunits (Hirst *et al*. 2011; Hirst *et al*. 2013), and interact with the adaptor protein complex AP-5 (Renvoisé *et al*. 2014; Słabicki *et al*. 2010; Hirst *et al*. 2011), by binding of the SPG11 N-terminus to AP-5 (Hirst *et al*. 2013). While this is superficially similar to the recruitment of clathrin to budding vesicles by adaptors, knockdown of *AP5z1* has in fact no effect on localization of SPG11 or SPG15, and knockdown of either SPG11 or SPG15 disrupts AP-5 localization (Hirst *et al*. 2013). Loss of SPG11, SPG15 or the AP-5 subunit AP5z1 causes accumulation of the mannose 6-phosphate receptor (CIMPR) in early endosomes (Hirst *et al*. 2013), and proteomic analyses of of AP5z1 knockout cells suggest a defect in transport from late endosomes to the Golgi (Hirst *et al*. 2018). Therefore, SPG11, SPG15 and AP-5 appear to share trafficking roles that may be disrupted in HSP.

SPG11 must also have AP-5-independent roles, which may account for disease phenotypes like ALS, PD or CMT, not so far found in human AP-5 mutant genotypes. Mutations in *AP5*z*1* are associated with HSP, although protein null alleles appear in general to have later onset (Hirst *et al*. 2016; Słabicki *et al*. 2010) than *SPG11* or *SPG15* (Pensato *et al*. 2014; Boukhris *et al*. 2008; Stevanin *et al*. 2007). Further, *Drosophila* contains both SPG11 and SPG15 homologs, but lacks AP-5 (Hirst *et al*. 2011). Such an AP-5-independent role appears to be in autophagic lysosome reformation (ALR) (Chang *et al*. 2014; Varga *et al*. 2015), which replenishes free lysosomes after periods of increased autophagy (Yu *et al*. 2010). Loss of SPG11 or SPG15 results in enlarged or excess autolysosomes, and loss of Lamp1-positive tubules on autolysosomes, thought to be an intermediate in ALR (Chang *et al*. 2014; Vantaggiato *et al*. 2013; Varga *et al*. 2015). Levels of LC3-II, an indicator of autophagy flux (Mauvezin *et al*. 2014), increase, but not Lamp1, suggesting aberrant lysosome-dependent clearance of autophagosomes (Chang *et al*. 2014; Varga *et al*. 2015). *Spg11* knock-out mice display fewer free lysosomes in Purkinje cell somata (Varga *et al*. 2015), consistent with impaired ALR *in vivo*. Mutant *SPG15* patient fibroblasts accumulate autophagy substrate p62 (Vantaggiato *et al*. 2013), and *SPG11* knock-out Purkinje cells show increased p62 in Lamp1-positive compartments, suggesting that autolysosomes are not completely functional (Varga *et al*. 2015). The above findings suggest a key role for SPG11 in ALR, and that ensuing accumulation of undegraded material in the absence of SPG11 may be a cause of neuronal degeneration (Varga *et al*. 2015).

However, the molecular role of SPG11 in autophagic trafficking, the steps it affects, and the nature of its AP-5-independent function are not clear. To identify potential roles for SPG11 in this process, we undertook bioinformatic analyses of the protein. We found that the presence or absence of the N-terminal region, found in most metazoan lineages, correlates with the presence or absence of the AP-5 adaptor complex, suggesting that the main function of the N-terminal region is to interact with AP-5. We also found evidence that the spatacsin_C domain at the C-terminus of SPG11 is homologous to the Vps16_C domain, and identified binding partners that might provide some common mechanisms for CMT pathology between SPG11 and aminoacyl-tRNA synthetases.

## Materials and Methods

### Amino acid alignments and protein function prediction

Multiple alignments of Spg11 sequences only were performed using Kalign multiple alignment, with gap open penalty of 13 (http://www.ebi.ac.uk/Tools/msa/kalign/) (Lassmann and Sonnhammer 2005), or MUSCLE multiple alignment (http://www.ebi.ac.uk/Tools/msa/muscle/) (Edgar 2004). Distant homologs were identified and aligned using HHpred (Zimmermann *et al*. 2018) on the MPI bioinformatic server (https://toolkit.tuebingen.mpg.de/hhpred). Tertiary structure prediction by homology was performed by submitting a multiple sequence alignment generated on the HHpred site to Modeller (Webb and Sali 2016). Interaction maps and GO and KEGG term enrichment were performed by submitting the list of *Drosophila* genes to String (http://string-db.org)

### Immunoprecipitation and mass spectrometry of GFP-tagged Spg11 C-terminus

We used Gateway recombination cloning technology (Invitrogen, UK) to generate a GFP::Spg11-C-term (nucleotides 3051-4094 of *Drosophila Spg11* gene). The entry and destination vectors were pDONR211 (Thermo Fisher, 12536017), Act5-Nterm GFP tag (driven by Actin5 promoter, pAGW plasmid, https://emb.carnegiescience.edu/drosophila-gateway-vector-collection) and PMT-pcBlast-Nterm GFP tag (CuSO4 Inducible promoter, Thermo Fisher V413020).

The C-terminal region of Spg11 (amino acid residues 1468-1815) was amplified using Phusion Flash High-Fidelity polymerase (Thermo Fisher F548L) and extracted from electrophoresis gel using QIAquick Gel Extraction Kit (Qiagen 28704) according to manufacturer’s protocol. The C-terminus from codons 1468-1815 was amplified using primers Spg11_C_f1 (GGGGACAAGTTTGTACAAAAAAGCAGGCTTC CAC CTG TGT TTC GTG CAC GA; forward sequence annealing to codons 1468-1473, HLCFVH shown as triplets) and Spg11_C_r1 (GGGGACCACTTTGTACAAGAAAGCTGGGTC CTA CGT ATC TCC TGC TGT GTG GA; complementary sequence annealing to codons 1810-1815 HTAGDT and stop, shown as triplets); they also contained additional homology to pDONR 211 allowing homologous recombination and integration of Spg11 C-terminus. BP and LR reactions conducted according to the Gateway® recombination cloning technology manual, and transformed into DH5-alpha cells according to the manufacturers instructions (Subcloning efficiency, ThermoFisher 18265017). Clones were sequenced (GATC-Biotech) using M13f and M13r primers, to ensure sequences were correct.

### Transfection of *Drosophila* Mel-2 Cells

*Drosophila* Mel-2 cells (Schneider 1972) were grown at 25°C in standard sterile cell flasks in Express SFM medium (LifeTechnologies), containing penicillin streptomycin (ThermoFisher) and L-glutamine (ThermoFisher). All cell work was carried out in a sterile hood. Prior to transfection, cells were grown to high confluence and seeded in 2 ml Express SFM (LifeTechnologies) in a 6-well plate, then left for 2 hours. To transform Mel-2 cells with GFP-tagged Spg11 C-terminus, 3 μg of plasmid DNA was diluted in 100 μl ddH_2_O in a microfuge tube. A helper plasmid (0.5 μg) was added in parallel with Act5-Nterm GFP tag to confer Mel-2 cells with blasticidin resistance. Next, 15 μl of FuGene-HD transfection reagent (Promega E2311) was applied directly to the center of the microfuge tube, and mixed thoroughly by gently pipetting up and down, before incubation for 15 minutes to allow formation of transfection complexes. Transfection mixture was added to the cells slowly, and the cell culture was subsequently incubated at 25°C for 12 hours.

Expression of GFP-tagged Spg11 C-terminus was confirmed by confocal microscopy. Mel-2 cells were seeded on sterile glass cover slips for 20 minutes, before fixing in ice cold methanol for 10 minutes. When necessary, cells were exposed to Lysotracker® for 10 minutes prior to fixation. Cells were mounted on glass slides before imaging.

Transfected Mel-2 cells were selected using blasticidin (ThermoFisher, USA, A1113903) for 5 passages. Cells were passaged when confluence approached 80%. They were then expanded to 7×10^8^ in 300 ml of Express SFM medium, in large sterile conical flasks, shaken at 80 rpm at 25°C. Transgene expression was induced 24 hours prior to protein extraction by adding 500 µM CuSO_4_. The PMT-pcBlast-Nterm GFP tag, lacking Spg11, was used as a control.

### *Drosophila* Mel-2 cell immunoprecipitation

*Drosophila* Mel-2 cells were transferred into 50 ml falcon tubes and pelleted at 5000 rpm, before resuspending cells in 8 ml of RIPA buffer (Sigma R0278), containing protease inhibitors (Sigma). Cells were lysed on ice using electronic homogenizer, 5 times, each for 30 seconds, with 30 seconds cool down time in between. Lysed cells were centrifuged for 30 minutes at 10,000 rpm at 4°C, before supernatant was collected. For immunoprecipitation, ChromoTek GFP-Trap A (ChromoTek gta-10, Germany) was used according to manufacturer’s instructions. GFP-tagged Spg11 C-terminus-bound beads were sent to for mass spectrometry analysis at the Mass Spectrometry Laboratory, Institute of Biochemistry and Biophysics, PAS. Poland.

### Data reporting

No institutional ethical review was required. The study was not pre-registered. Randomization, blinding, and sample size calculations were not applicable. No animals were used. The study was exploratory. Statistical calculations were all carried out by the bioinformatic software described. Clones can be provided on request, or generated from the information provided in the paper.

## Results

### N-terminal AP-5-binding region of SPG11 is absent in insect lineages

We used bioinform-atic tools to identify potential functional domains within SPG11. Multiple alignments show extensive conservation along the length of SPG11 proteins, but with no identifiable domains of known function (Hirst *et al*. 2011; Hirst *et al*. 2013). The best characterized molecular function is a predicted beta-propeller-like fold in the N-terminal 500 amino-acid residues of human SPG11, which interacts with the AP-5 adaptor (Hirst *et al*. 2013). Multiple alignment (Lassmann and Sonnhammer 2005) identified this region in all deuterostomes aligned (Fig. 1), and in many protostomes, including arachnids, molluscs, annelids and brachiopods. However, *Drosophila* and other insects lack approximately 650 amino acids from the N-terminus of SPG11 (known as CG13531 in *Drosophila*), which contains the AP-5-binding domain (Fig. 1).

**Fig. 1.**
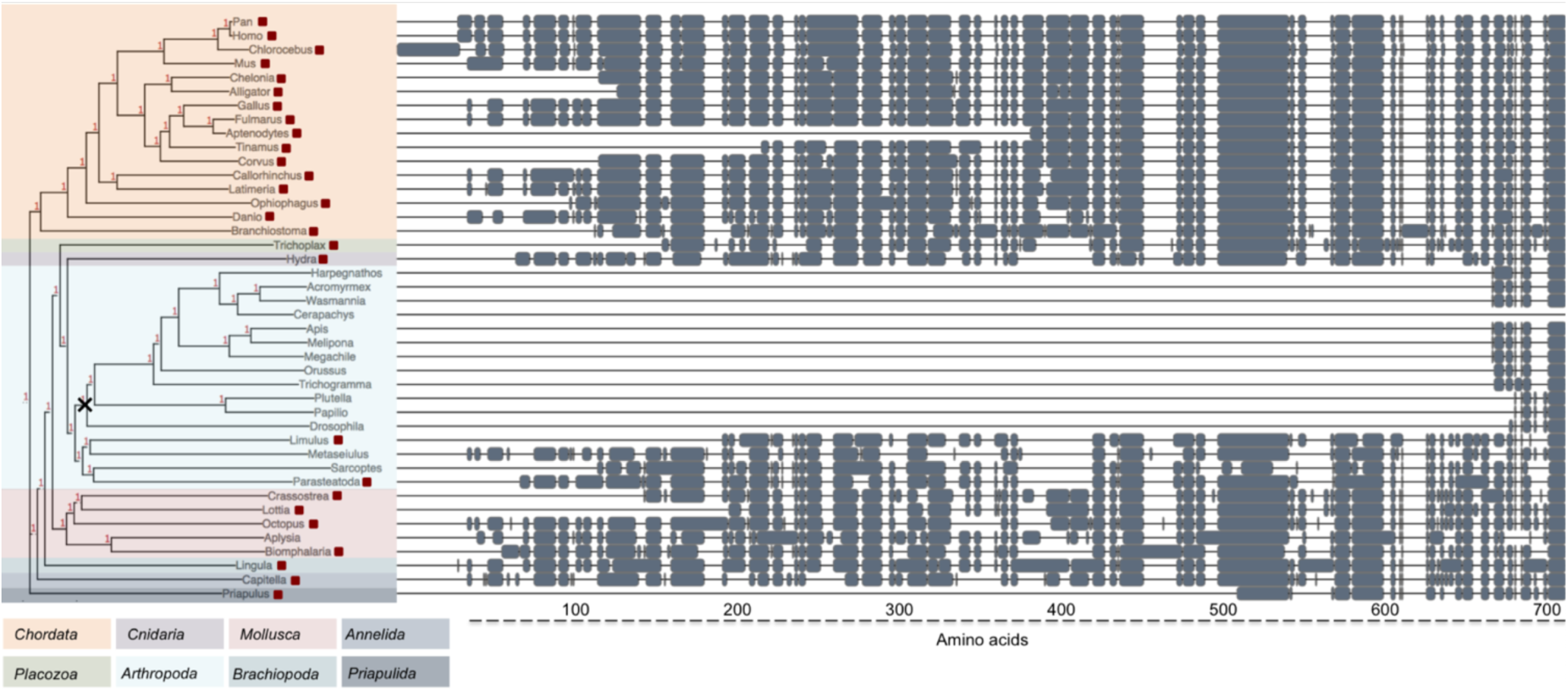
SPG11 phylogenetic tree across a range of metazoans. Left: Phylogenetic tree of SPG11 amino acid sequence, made using Kalign multiple alignment (Lassmann and Sonnhammer 2005). Organisms containing AP-5 (Figs S1-S4) are shown by red squares. Shaded regions denote different phyla. Right: A condensed overview of aligned SPG11 N-terminal regions, created using online tree viewer (http://etetoolkit.org/treeview/). Predicted loss of N-terminal region occurs prior to the divergence of insects (cross).

AP-5 is an ancient adaptor complex, but has frequently been lost during eukaryote evolution. To test whether its loss correlates with loss of the SPG11 N-terminus, we tested a range of metazoans for the presence of any of the four AP-5 subunits (Hirst *et al*. 2011). AP-5 is present in basal animal phyla such as *Cnidaria*, the sponge *Amphimedon*, and the Placozoan *Trichoplax*; it is also found widely in deuterostomes (Figs S1-S4). In protostomes, it is found in molluscs (*Octopus, Lottia, Crassostrea, Biomphalaria*), the brachiopod *Lingula*, the annelid *Capitella*, the priapulid worm *Priapulus*, and some arthropods (*Limulus, Parasteatoda*), but not in insects. Most organisms that lack the SPG11 N-terminal region (Figure 1) also lack AP-5 (Figs S1-S4), suggesting that the primary role of the N-terminus of SPG11 is to bind AP-5.

In a few species, we found an SPG11 N-terminal domain, but not AP-5: the arthropods *Sarcoptes* and *Metaseiulus*, and the mollusc *Aplysia* (Figure 1). However, the arthropods most closely related to *Sarcoptes* and *Metaseiulus* (*Limulus, Parasteatoda*), and other molluscs (*Octopus, Lottia*), possess both AP-5 and the SPG11 N-terminus, suggesting either that AP-5 is present in *Sarcoptes, Metaseiulus* and *Aplysia* but not yet annotated, or these species have lost AP-5 relatively recently, but not yet the SPG11 N-terminus.

### SPG11 C-terminus has predicted homology with the Vps33-binding domain of Vps16

Human and *Drosophila* SPG11 share 16% identity across the entire protein sequence, with higher identity (31%) across a C-terminal region of around 300 amino acid residues (Fig. 2A). This region is designated as a Spatacsin_C domain (Pfam 14649; Interpro IPR028107), but its function or structure is unknown. To identify domains homologous to this region of SPG11, we used *Drosophila* and human SPG11 C-terminal domain sequences as queries in a HHpred search; this search can identify distant homologies between proteins using the Structural Classification of Proteins database (SCOP) (Söding *et al*. 2005), and previously predicted a beta-propeller-like region in the N-terminal AP-5-binding domain (Hirst *et al*. 2013). It also predicts homology of some regions of *Drosophila* SPG11 to coatomer subunits (Figure 2B), as previously predicted for human SPG11. At the C-terminus, HHpred predicted homology across most of the C-terminal 300 amino acid residues, to PDB crystal structure 4KMO (DOI 10.2210/pdb4kmo/pdb), comprising the Vps33-Vps16 HOPS subcomplex (Figure 2B,C), which contains the C-terminal region of Vps16 (Vps16_C, Pfam ID: PF04840) and the SM protein Vps33 (Baker *et al*. 2013; Graham *et al*. 2013).

**Fig. 2.**
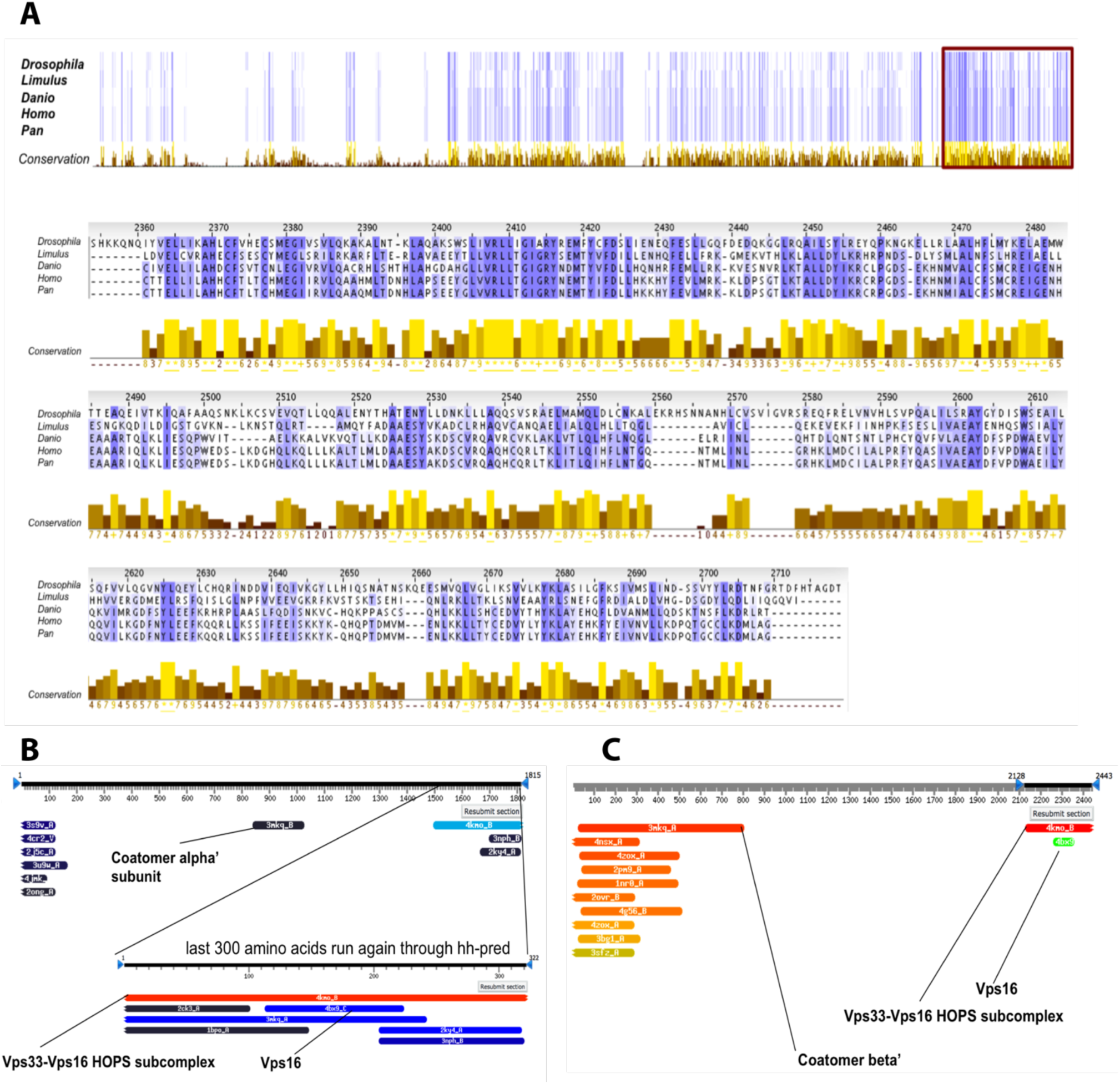
Sequence conservation across the length of Spg11. **A**. MUSCLE multiple alignment of SPG11 from *D. melanogaster, L. polyphemus, D. rerio, H. sapiens* and *P. troglodytes*, shown in Jalview. Top panel: blue shaded regions show regions of amino acid identity between organisms; intensity of blue corresponds with level of identity. Yellow bars at the bottom denote conservation level between organisms for individual amino acid positions. Red box indicates SPG11 C-terminal region with high conservation. This region is expanded in the lower panels to show conservation and alignment of amino acid sequences. **B:** HHpred of *Drosophila* SPG11 predicts coatomer homology in the central region of the proteins, and homology to Vps33-Vps16 HOPS subcomplex and Vps16 in the C-terminal region. **C:** HHpred analysis of human SPG11 predicts extended homology to coatomer beta’ subunit at the N-terminus and to Vps33-Vps16 HOPS subcomplex and Vps16 at the C-terminus.

To verify the sequence conservation of the SPG11 C-terminus to the Vps33-Vps16 subcomplex, we used six phylogenetically diverse SPG11 C-terminal domains as a single query in a HHpred alignment. After other Spg11 C-termini, the next most closely related sequences were the C-terminal domains of the Vps16a and Vps16b families (Supplementary Fig. 5). A multiple sequence alignment in HHpred using a small number of Spg11, Vps16A and Vps16B proteins, found extensive similarity between all three families of proteins, with identical or conserved amino acid residues at many positions (Figure 3A). However, many known Vps33-binding residues of the Vps16_C domain of Vps16 (Baker *et al*. 2013; Graham *et al*. 2013) were either missing or not conserved in SPG11 (Figure 3A). Together these results support homology of the SPG11 C-terminal domain to Vps16_C, but do not strongly support binding to Vps33.

**Fig. 3.**
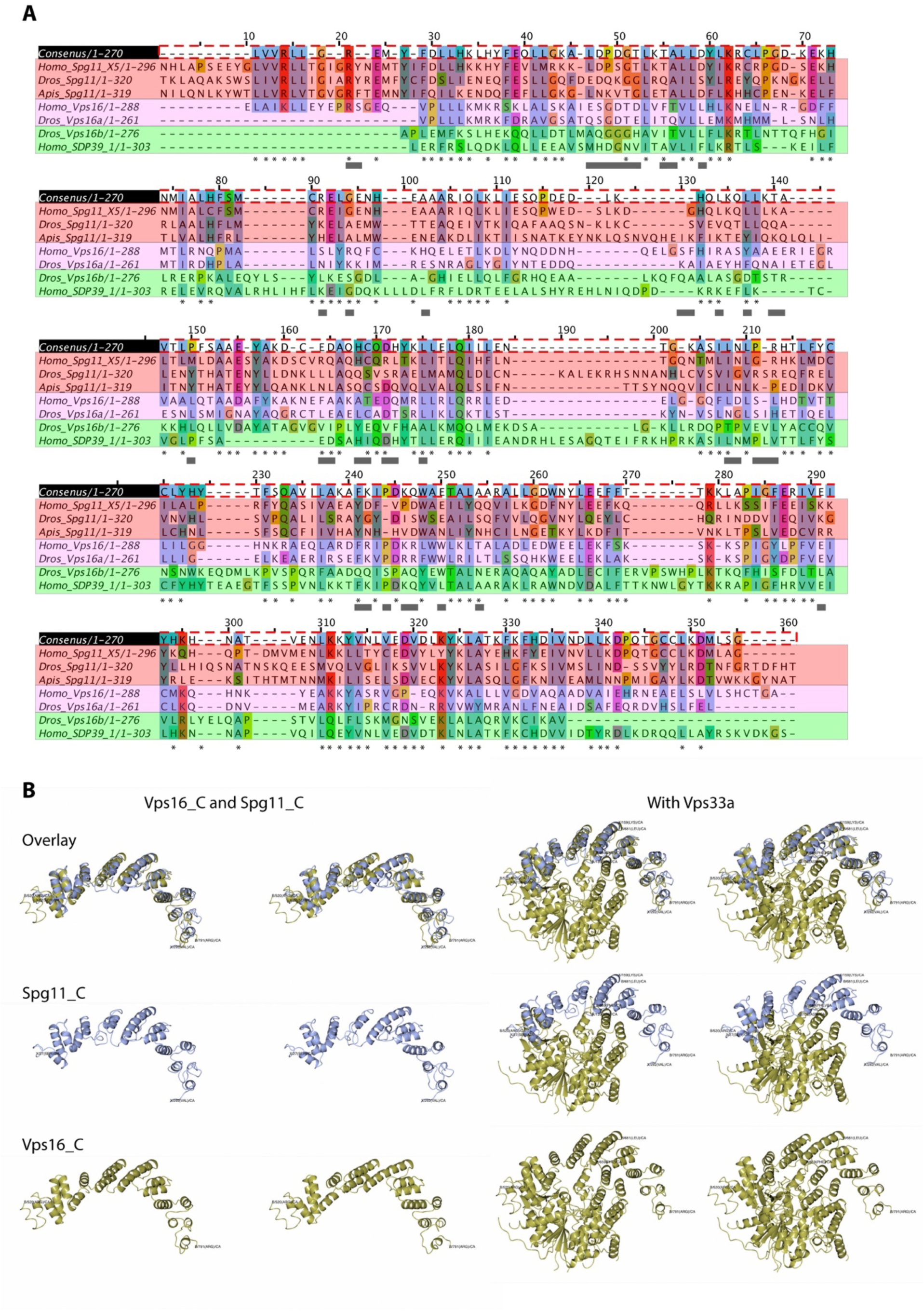
SPG11 C-terminus shows conservation with Vps16_C domains. **A**. Multiple alignment of Spg11 C-terminus with sequences from two Vps16A and two Vps16B subfamily members, grouped by color. We generated a multiple sequence alignment in HHpred (Zimmermann *et al*. 2018), exported to JALview, and made minor adjustments manually. Asterisks below alignment show residues that mostly conserved, or show only conservative changes, between the Spg11 sequences and at least one of the Vps16 sequences. Gray boxes below alignment denote Vps33-binding residues in Vps16a (Baker *et al*. 2013; Graham *et al*. 2013). **B**. Tertiary structure prediction of SPG11_C was performed by forwarding a HHpred multiple sequence alignment of six Spg11_C domains from the HHpred site to Modeller (Webb and Sali 2016), to align with the tertiary structure of Vps16_C (4KMO:B, https://www.rcsb.org/). Stereo pairs show actual structure of Vps16_C and predicted structure of Spg11_C, either alone (left pairs, overlaid in top row, singly in lower rows), or bound to Vps33a (right pairs).

### SPG11 C-terminus predicted tertiary structure similar to Vps16_C

The Vps16_C domain forms an alpha-solenoid with 17 alpha-helices; it binds to all 3 domains of Vps33, arching around Vps33, its N-terminal region lying in a groove between Vps33 domains 1 and 2 (Baker *et al*. 2013; Graham *et al*. 2013). We assessed whether the SPG11 C-terminus might adopt a similar structure to Vps16_C, using Modeller tertiary structure prediction (Webb and Sali 2016). To give more weight to conserved residues for modeling, we tested the multiple alignment of six Spg11 sequences (Supplementary Fig. S5) for predicted similarity to the tertiary structure of Vps16_C (4KMO:B, https://www.rcsb.org/). This analysis predicted an extensive alpha-helical structure for the Spg11 C-terminus, with the majority of helices at positions compatible with those in Vps16_C (Fig. 3B). The potential of the Spg11 C-terminus to adopt a tertiary structure like Vps16_C further supports a model of its origin as a distant homolog of Vps16_C.

### GFP-tagged SGP11 C-terminus overlaps with acidic compartments

To investigate whether the SPG11 C-terminus localized to endo/auto/lysosomal compartments like Vps16 (Kim *et al*. 2001; Schröder *et al*. 2007; Wartosch *et al*. 2015) and Spg11 (Hirst *et al*. 2012; Murmu *et al*. 2011; Varga *et al*. 2015), we expressed GFP::Spg11-C (amino acid residues 1468-1815) in *Drosophila* Mel-2 cells. GFP::Spg11-C localized widely through the cytosol, but was often concentrated in puncta that overlapped with acidic LysoTracker®-positive compartments in many cells (Figure 4), showing that much of it localized to endo/auto/lysosomal compartments.

**Fig. 4.**
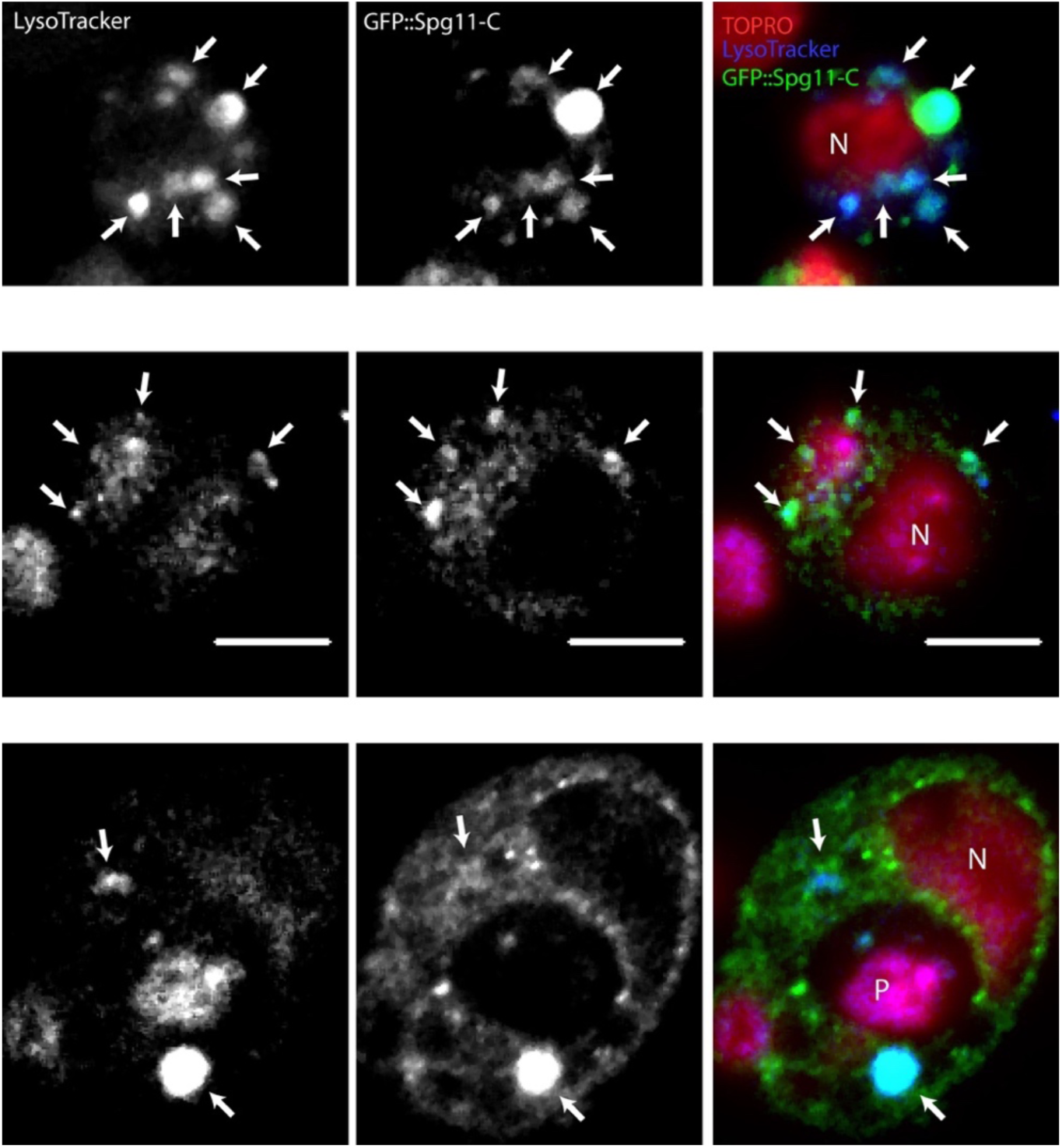
Localisation of GFP::Spg11-C in *Drosophila* Dmel-2 cells. Single sections of three different transfected cells, showing overlap of many GFP::Spg11-C puncta with LysoTracker® -positive acidic compartments (examples shown by arrows). Nuclei (N) are stained lightly with TOPRO-3. One likely phaogosome (P) is visible. Scale bar = 5 μm.

### Identification of SPG11 C-terminus binding partners by mass spectrometry

To test further the functionality of the SPG11 C-terminus, we conducted mass spectrometry analysis on *Drosophila* GFP::Spg11-C expressed in Mel-2 cells. We established two cell lines: one with GFP::Spg11-C driven by a constitutively active promoter (Act5-GFP::Spg11-C) and one by a CuSO_4_-inducible promoter (PMT-GFP::Spg11-C). A Mel-2 cell line expressing GFP (inducible by CuSO_4_) was used as a negative control.

Mass spectrometry analysis of purified GFP::Spg11-C from CuSO_4_-induced cells identified 898 proteins not found in the GFP negative control, whereas analysis of constitutively expressed GFP::Spg11-C identified 231 proteins (Fig. 5A); the top 100 hits from each analysis are listed in Tables 1 and 2. The CuSO_4_-induced cells generated more protein hits that showed higher scores than in the control cell line, compared to the constitutive cells (Figure 5B), suggesting that GFP::Spg11-C was more highly expressed in CuSO_4_-inducible than in constitutive cells. 128 proteins were detected in both lines (Fig. 5A; Table 3), providing increased confidence in the specificity of these interactors, although this overlapping list may be missing some genuine interactors identified in the CuSO_4_-induced cells.

**Table 1.**
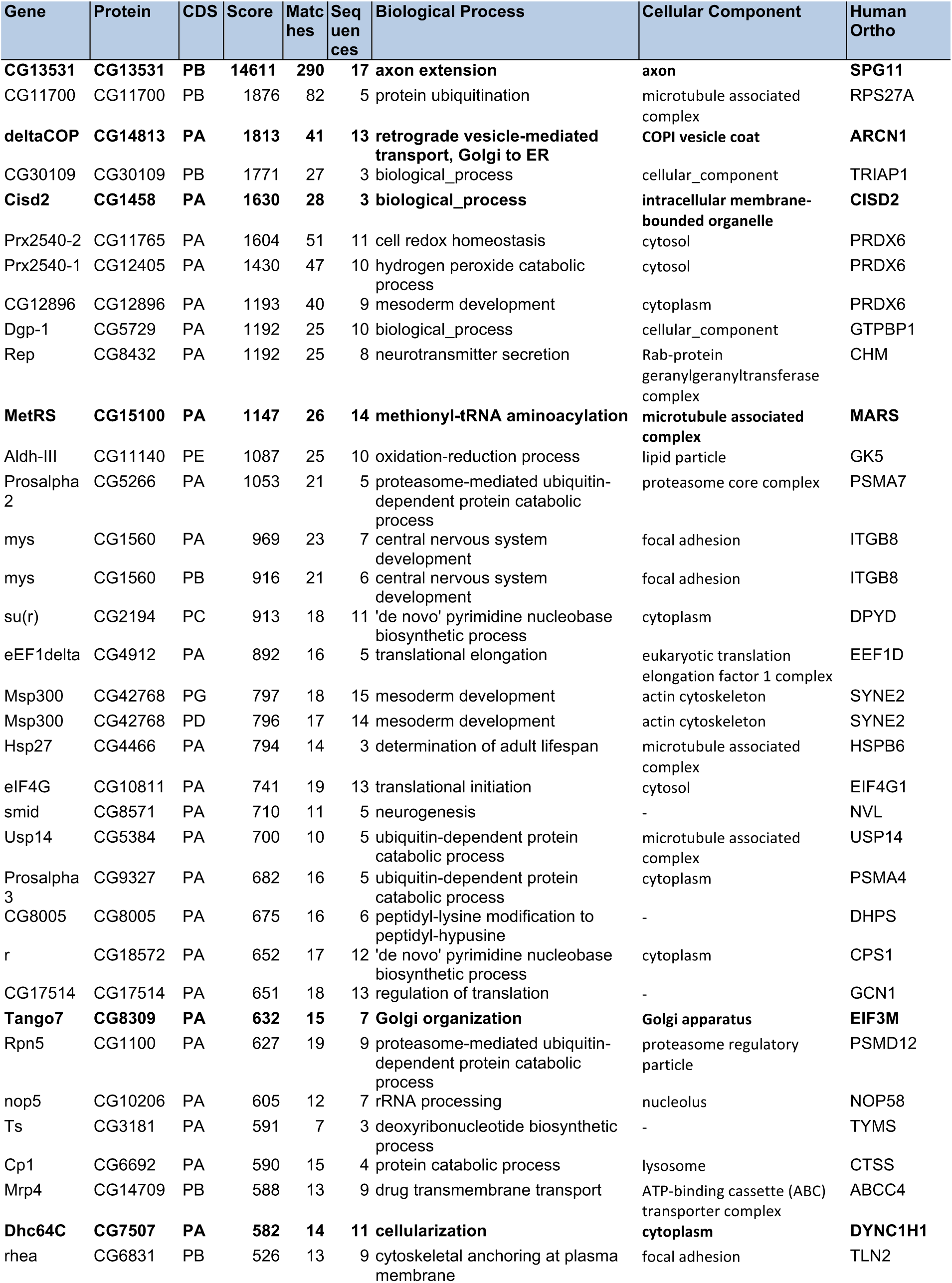
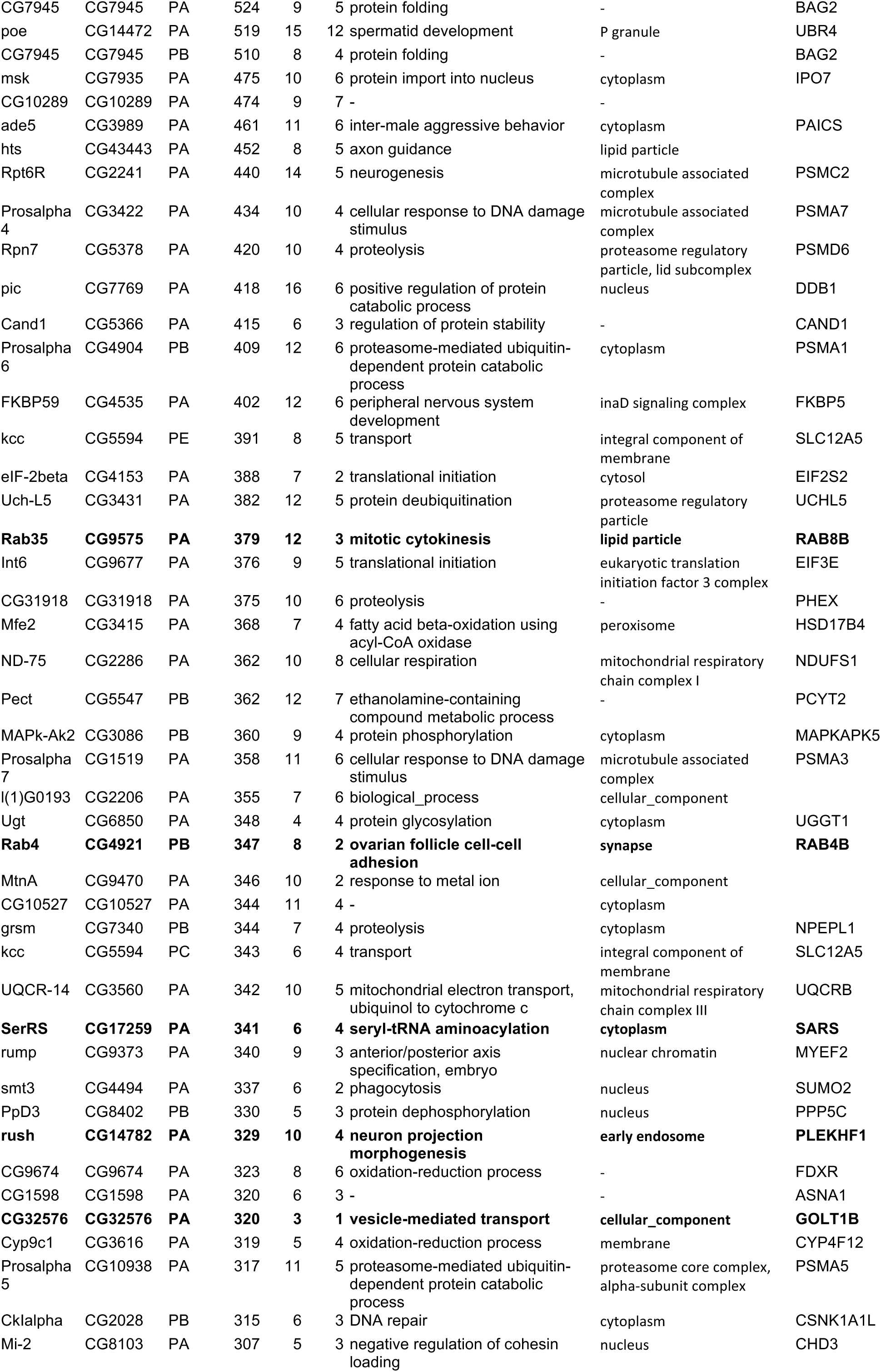
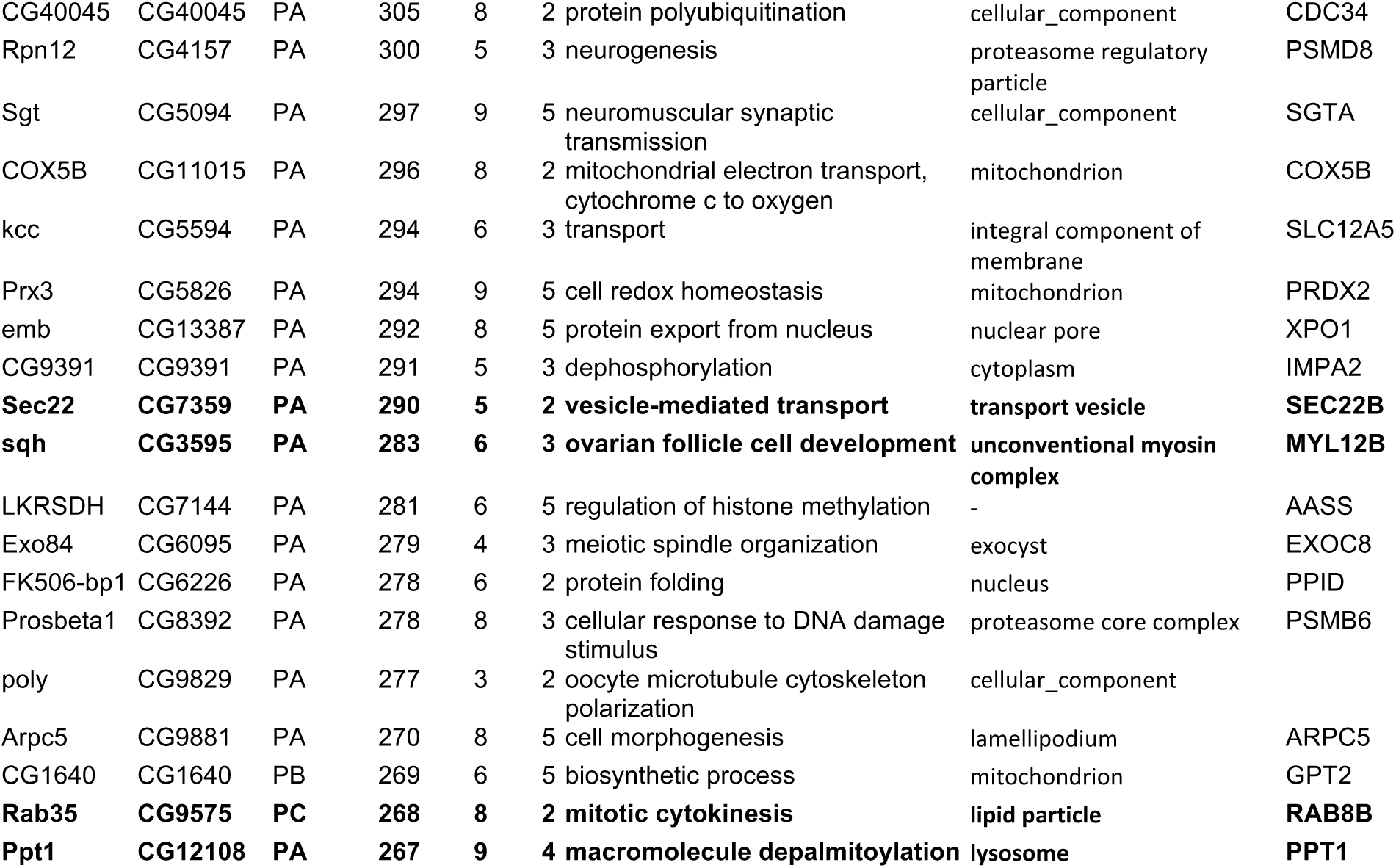
Top 100 unique hits found in mass spectrometry output from GFP::Spg11-C purification from PMT-GFP::Spg11-C. Biological process and human ortholog (Human Ortho) data are from Flybase (www.flybase.org). Some of the hits discussed in the text are in bold.

**Table 2.**
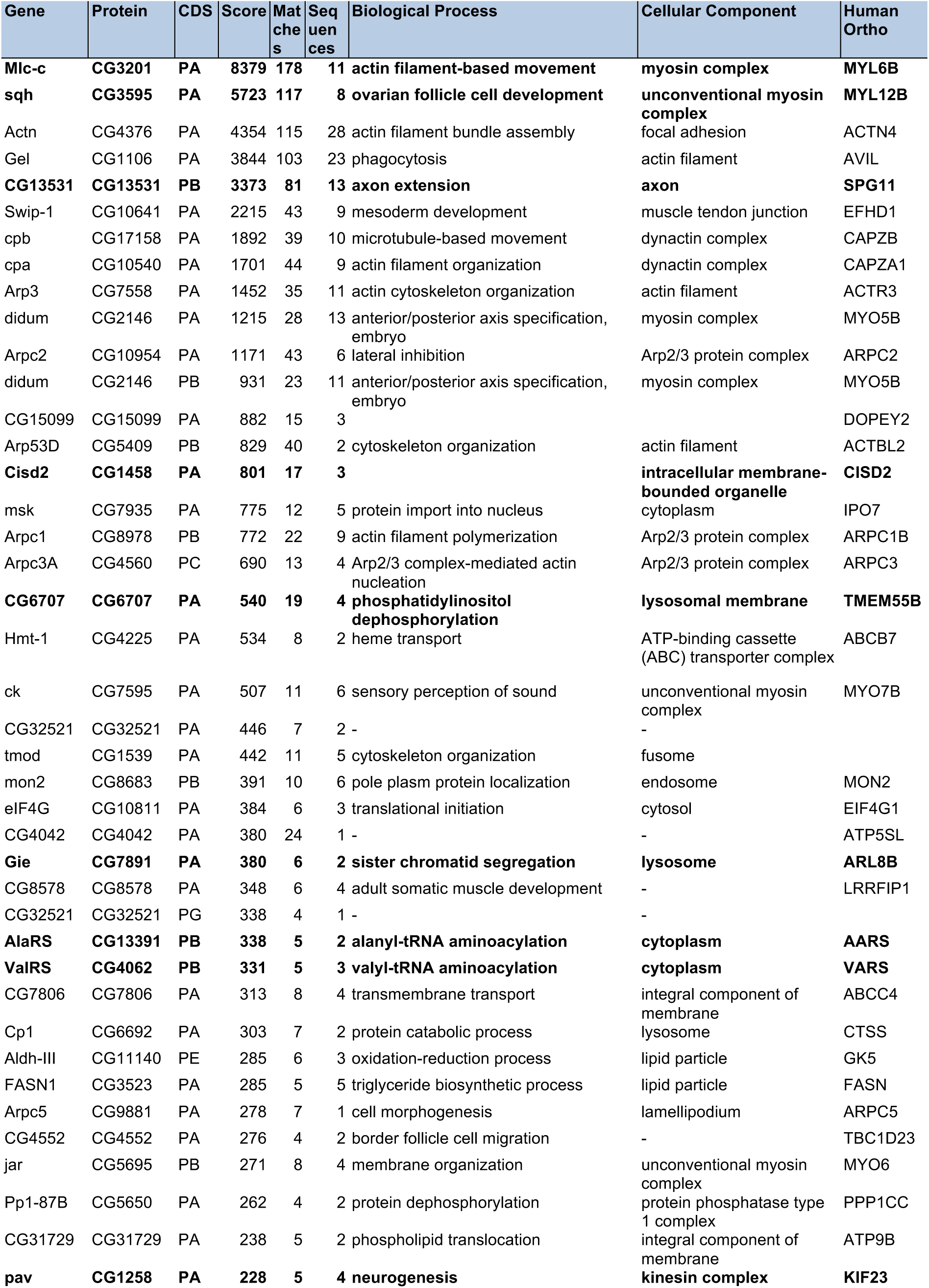
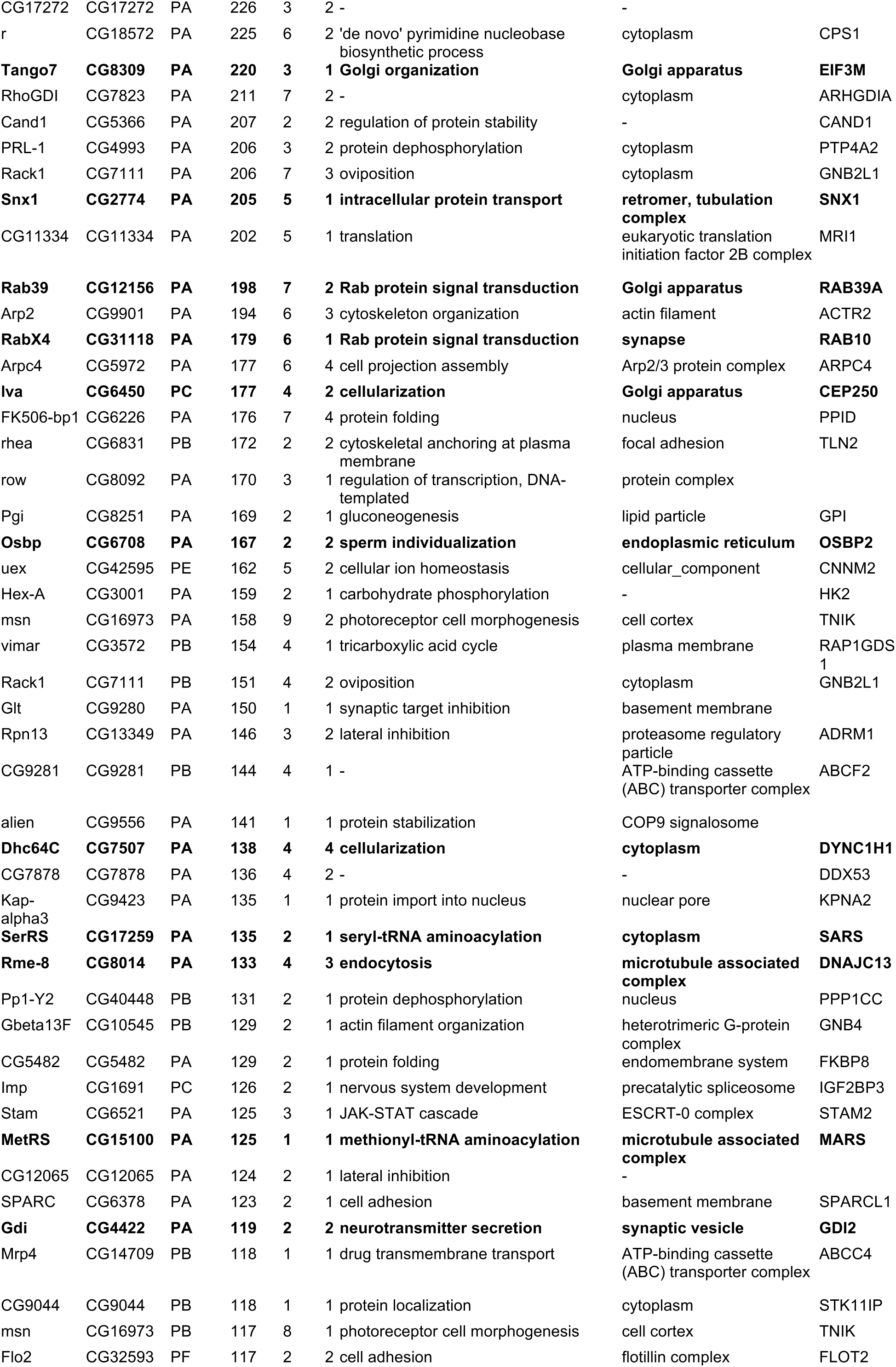
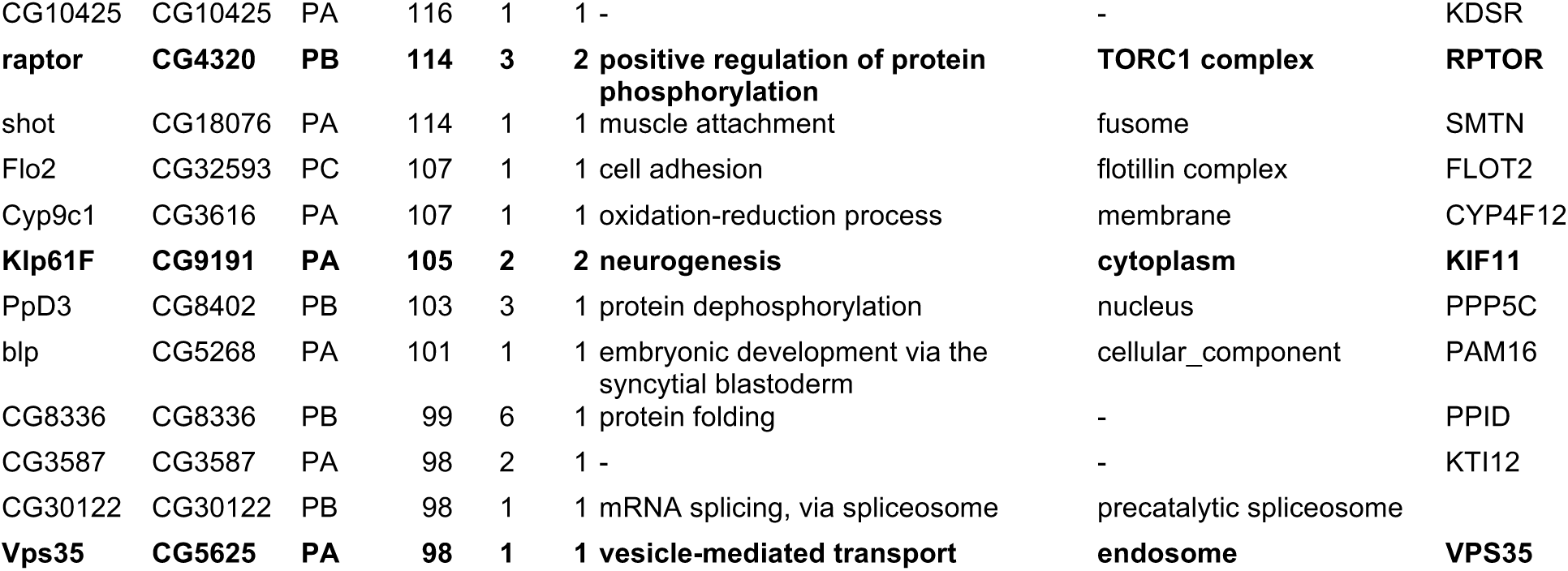
Top 100 unique hits found in mass spectrometry output from GFP::Spg11-C purification from Act-Spg11-C::GFP. Biological process and human ortholog (Human Ortho) data from Flybase (www.flybase.org). Some of the hits discussed in the text are in bold.

**Table 3.** Protein hits identified from GFP::Spg11-C purification from both Act5-GFP::Spg11-C and PMT-GFP::Spg11-C. Proteins are ranked by highest score present in either of the two lines. Proteins mentioned in the text are highlighted. Human orthologue (Human Orth) data are from Flybase (www.flybase.org). Some of the hits discussed in the text are in bold.

| # | Protein | Gene | Human Orth | # | Protein | Gene | Human Orth | # | Protein | Gene | Human Orth |
| --- | --- | --- | --- | --- | --- | --- | --- | --- | --- | --- | --- |
| 1 | **CG13531** | **CG13531** | **SPG11** | 51 | **CG15811** | **Rop** | **STXBP3** | 101 | CG1828 | *dre4* | SUPT16H |
| 2 | CG5409 | *Arp53D* | ACTBL2 | 52 | CG13849 | *Nop56* | NOP58 | 102 | CG7207 | *cert* | COL4A3BP |
| 3 | CG2241 | *Rpt6R* | PSMC2 | 53 | CG10990 | *Pdcd4* | PDCD4 | 103 | CG6020 | *ND-39* | ENS<br>G00000255639 |
| 4 | CG7878 | *CG7878* | DDX53 | 54 | CG5650 | *Pp1-87B* | PPP1CC | 104 | CG5336 | *Ced-12* | ELMO3 |
| 5 | CG1458 | *Cisd2* | CISD2 | 55 | CG3523 | *FASN1* | FASN | 105 | CG1484 | *flil* | FLII |
| 6 | CG11765 | *Prx2540-2* | PRDX6 | 56 | CG5504 | *l(2)tid* | DNAJA3 | 106 | CG4993 | *PRL-1* | PTP4A2 |
| 7 | **CG12156** | **Rab39** | **RAB39A** | 57 | CG3991 | *TppII* | TPP2 | 107 | CG4042 | *CG4042* | ATP5SL |
| 8 | **CG31118** | **RabX4** | **RAB10** | 58 | CG12400 | *ND-B14.5B* |  | 108 | CG43398 | *scrib* | SCRIB |
| 9 | CG5729 | *Dgp-1* | GTPBP1 | 59 | CG11804 | *ced-6* | GULP1 | 109 | **CG2774** | **Snx1** | **SNX1** |
| 10 | **CG15100** | **MetRS** | **MARS** | 60 | **CG3201** | **Mlc-c** | **MYL6B** | 110 | CG6963 | *gish* | CSNK1G3 |
| 11 | CG11140 | *Aldh-III* | GK5 | 61 | CG10539 | *S6k* | RPS6KA3 | 111 | CG12202 | *NAA15-16* | NAA15 |
| 12 | CG15871 | *mRpL38* | MRPL38 | 62 | **CG5625** | **Vps35** | **VPS35** | 112 | CG5482 | *CG5482* | FKBP8 |
| 13 | CG10811 | *eIF4G* | EIF4G1 | 63 | CG4931 | *Sra-1* | CYFIP1 | 113 | **CG6708** | **Osbp** | **OSBP2** |
| 14 | CG18572 | *r* | CPS1 | 64 | CG1987 | *Rbp1-like* | SRSF3 | 114 | CG17158 | *cpb* | CAPZB |
| 15 | CG17514 | *CG17514* | GCN1 | 65 | CG18076 | *shot* | SMTN | 115 | CG9186 | *CG9186* | LDHA |
| 16 | **CG8309** | **Tango7** | **EIF3M** | 66 | CG32056 | *scramb1* | PLSCR1 | 116 | **CG6450** | **Iva** | **CEP250** |
| 17 | CG1100 | *Rpn5* | PSMD12 | 67 | CG30122 | *CG30122* |  | 117 | CG8578 | *CG8578* | LRRFIP1 |
| 18 | CG10206 | *nop5* | NOP58 | 68 | CG7558 | *Arp3* | ACTR3 | 118 | CG7421 | *Nopp140* | NOLC1 |
| 19 | CG6692 | *Cp1* | CTSS | 69 | CG7324 | *CG7324* | TBC1D9 | 119 | CG8207 | *CG8207* | GMPPA |
| 20 | CG14709 | *Mrp4* | ABCC4 | 70 | **CG4062** | **ValRS** | **VARS** | 120 | CG30159 | *CG30159* |  |
| 21 | **CG7507** | **Dhc64C** | **DYNC1H1** | 71 | CG7111 | *Rack1* | GNB2L1 | 121 | CG7985 | *CG7985* | HEXDC |
| 22 | CG6831 | *rhea* | TLN2 | 72 | **CG10506** | **GlnRS** | **QARS** | 122 | CG43947 | *PsGEF* |  |
| 23 | CG7935 | *msk* | IPO7 | 73 | CG8261 | *Ggamma1* | GNG10 | 123 | CG7035 | *Cbp80* | NCBP1 |
| 24 | CG43443 | *hts* |  | 74 | CG3585 | *Rbcn-3A* | DMXL1 | 124 | CG5268 | *blp* | PAM16 |
| 25 | CG5366 | *Cand1* | CAND1 | 75 | CG32138 | *Frl* | DAAM2 | 125 | CG8989 | *His3.3B* | HIST1H3B |
| 26 | CG5594 | *kcc* | SLC12A5 | 76 | CG4422 | *Gdi* | GDI2 | 126 | CG7538 | *Mcm2* | MCM2 |
| 27 | CG3431 | *Uch-L5* | UCHL5 | 77 | CG3299 | *Vinc* | VCL | 127 | **CG6707** | **CG6707** | **TMEM55B** |
| 28 | **CG17259** | **SerRS** | **SARS** | 78 | CG9423 | *Kap-alpha3* | KPNA2 | 128 | **CG4320** | **raptor** | **RPTOR** |
| 29 | CG8402 | *PpD3* | PPP5C | 79 | CG10545 | *Gbeta13F* | GNB4 |  |  |  |  |
| 30 | CG3616 | *Cyp9c1* | CYP4F12 | 80 | CG10913 | *Spn55B* | SERPINB2 |  |  |  |  |
| 31 | CG5094 | *Sgt* | SGTA | 81 | CG4265 | *Uch* | UCHL3 |  |  |  |  |
| 32 | CG11015 | *COX5B* | COX5B | 82 | CG3001 | *Hex-A* | HK2 |  |  |  |  |
| 33 | **CG3595** | **sqh** | **MYL12B** | 83 | **CG8014** | **Rme-8** | **DNAJC13** |  |  |  |  |
| 34 | CG6226 | *FK506-bp1* | PPID | 84 | CG8683 | *mon2* | MON2 |  |  |  |  |
| 35 | CG9881 | *Arpc5* | ARPC5 | 85 | CG17950 | *HmgD* | SSRP1 |  |  |  |  |
| 36 | CG12108 | *Ppt1* | PPT1 | 86 | CG8284 | *Ubc4* | UBE2K |  |  |  |  |
| 37 | CG7619 | *Rpn10* | PIP5K1C | 87 | CG12918 | *sel* | CNPY1 |  |  |  |  |
| 38 | CG9281 | *CG9281* | ABCF2 | 88 | CG5252 | *Ranbp9* | IPO9 |  |  |  |  |
| 39 | CG10425 | *CG10425* | KDSR | 89 | CG8487 | *garz* | GBF1 |  |  |  |  |
| 40 | CG13349 | *Rpn13* | ADRM1 | 90 | CG12132 | *c11.1* | MROH7 |  |  |  |  |
| 41 | CG3226 | *CG3226* | CACYBP | 91 | CG12065 | *CG12065* |  |  |  |  |  |
| 42 | CG34373 | *Sarm* | SARM1 | 92 | CG4225 | *Hmt-1* | ABCB7 |  |  |  |  |
| 43 | CG16973 | *msn* | TNIK | 93 | CG10161 | *eIF-3p66* | EIF3D |  |  |  |  |
| 44 | CG8251 | *Pgi* | GPI | 94 | CG14616 | *l(1)G0196* | PPIP5K1 |  |  |  |  |
| 45 | CG9242 | *bur* | GMPS | 95 | CG42595 | *uex* | CNNM2 |  |  |  |  |
| 46 | **CG13391** | **AlaRS** | **AARS** | 96 | CG3572 | *vimar* | RAP1GDS1 |  |  |  |  |
| 47 | CG1691 | *Imp* | IGF2BP3 | 97 | CG11334 | *CG11334* | MRI1 |  |  |  |  |
| 48 | CG6783 | *fabp* | FABP2 | 98 | CG4560 | *Arpc3A* | ARPC3 |  |  |  |  |
| 49 | CG3593 | *r-l* | UMPS | 99 | CG7806 | *CG7806* | ABCC4 |  |  |  |  |
| 50 | **CG7891** | **Gie** | **ARL8B** | 100 | CG8715 | *lig* | UBAP2 |  |  |  |  |

GFP::Spg11-C was detected in both cell lines, but not in the GFP negative control line, confirming its expression, and the mass spectrometry protocol (Figure 5B); it was the highest scoring hit in the CuSO_4_-induced line, but not in the constitutively expressing cell line (Figure 5B). Reassuringly, the SPG11 binding partner, SPG15, was identified in the inducible cell line, although not in the constitutive one. To assess the potential biological roles of Spg11, we tested for enrichment of molecular function, biological process, and pathway terms among the interactors identified in both cell lines. Membrane traffic and organization terms were among those most enriched (Table 4; Fig. 6). We also noticed several actin-binding proteins, and motor proteins of the kinesin, dynein and myosin families (Tables 1-3, entries in bold; Fig. 6).

**Table 4.** Enrichment analysis for Spg11 C-terminal interactors identified in both constitutive and inducible interaction screens. Enrichment analysis is shown for human orthologs. GO terms with large gene sets (>2000) and smaller sets with less specific terms are omitted. Full lists are in Supplementary Information.

| GO-term | Molecular Function | Count in gene set | False Discovery Rate |
| --- | --- | --- | --- |
| GO:0008144 | drug binding | 32 of 1710 | 2.97E-06 |
| GO:0005524 | ATP binding | 29 of 1462 | 2.97E-06 |
| GO:0004812 | aminoacyl-tRNA ligase activity | 5 of 43 | 0.00032 |
| GO:0003723 | RNA binding | 17 of 850 | 0.00049 |
| GO:0140296 | general transcription initiation factor binding | 4 of 39 | 0.00300 |
| GO:0045182 | translation regulator activity | 6 of 124 | 0.00300 |
| GO:0003779 | actin binding | 10 of 413 | 0.00470 |
| GO-term | Biological Process |  |  |
| GO:0006897 | vesicle-mediated transport | 30 of 1699 | 0.00022 |
| GO:0016192 | regulated exocytosis | 17 of 691 | 0.00043 |
| GO:0051234 | neutrophil degranulation | 14 of 485 | 0.00060 |
| GO:0043604 | phagocytosis | 9 of 185 | 0.00063 |
| GO:0006810 | endocytosis | 14 of 510 | 0.00063 |
| GO:0045055 | translation | 12 of 362 | 0.00063 |
| GO:0043312 | cell activation involved in immune response | 15 of 620 | 0.00075 |
| GO:1901566 | secretion | 20 of 1070 | 0.00086 |
| GO:0006909 | membrane organization | 16 of 729 | 0.00110 |
| KEGG Pathway | Description |  |  |
| dme00970 | Aminoacyl-tRNA biosynthesis | 5 of 42 | 0.0031 |
| dme04144 | Endocytosis | 6 of 119 | 0.0236 |
| dme03050 | Proteasome | 4 of 51 | 0.0236 |
| dme02010 | ABC transporters | 3 of 22 | 0.0236 |

**Fig. 5.**
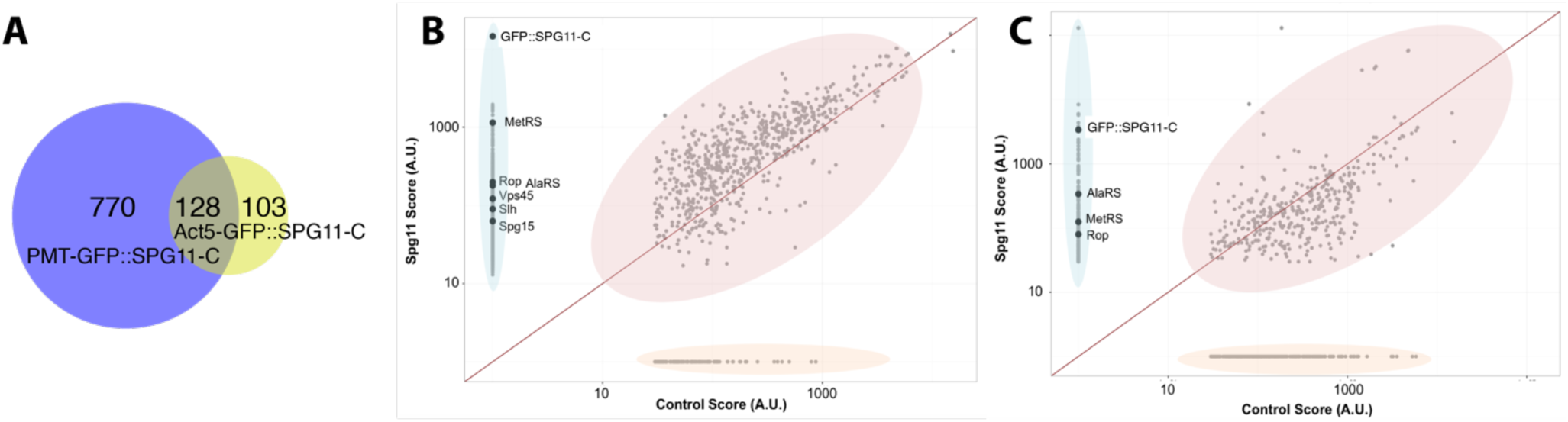
Binding partners of Spg11_C identified by mass spectrometry. **A**. Venn diagram of Spg11_C interactors identified by mass spectrometry of PMT-GFP::Spg11-C and Act5-GFP::Spg11-C cell lines. **B**,**C**. Mass spectrometry output for purified GFP::Spg11-C from the *Drosophila* Mel-2 cell line expressing PMT-GFP::Spg11-C (**B**) or Act5-GFP::Spg11-C (**C**), compared to control cells lacking GFP::Spg11-C. Graphs plot the percentage difference between test and control. **B** has more proteins with a higher score than the equivalent protein in the negative GFP control, indicated by the majority of the protein hits residing above the equivalence line (in red). However, graph C has fewer proteins above the equivalence line. Scores of GFP::Spg11-C, Rop, Vps45, Slh, MetRS (MARS), AlaRS (AARS) and Spg15 are highlighted.

**Fig. 6.**
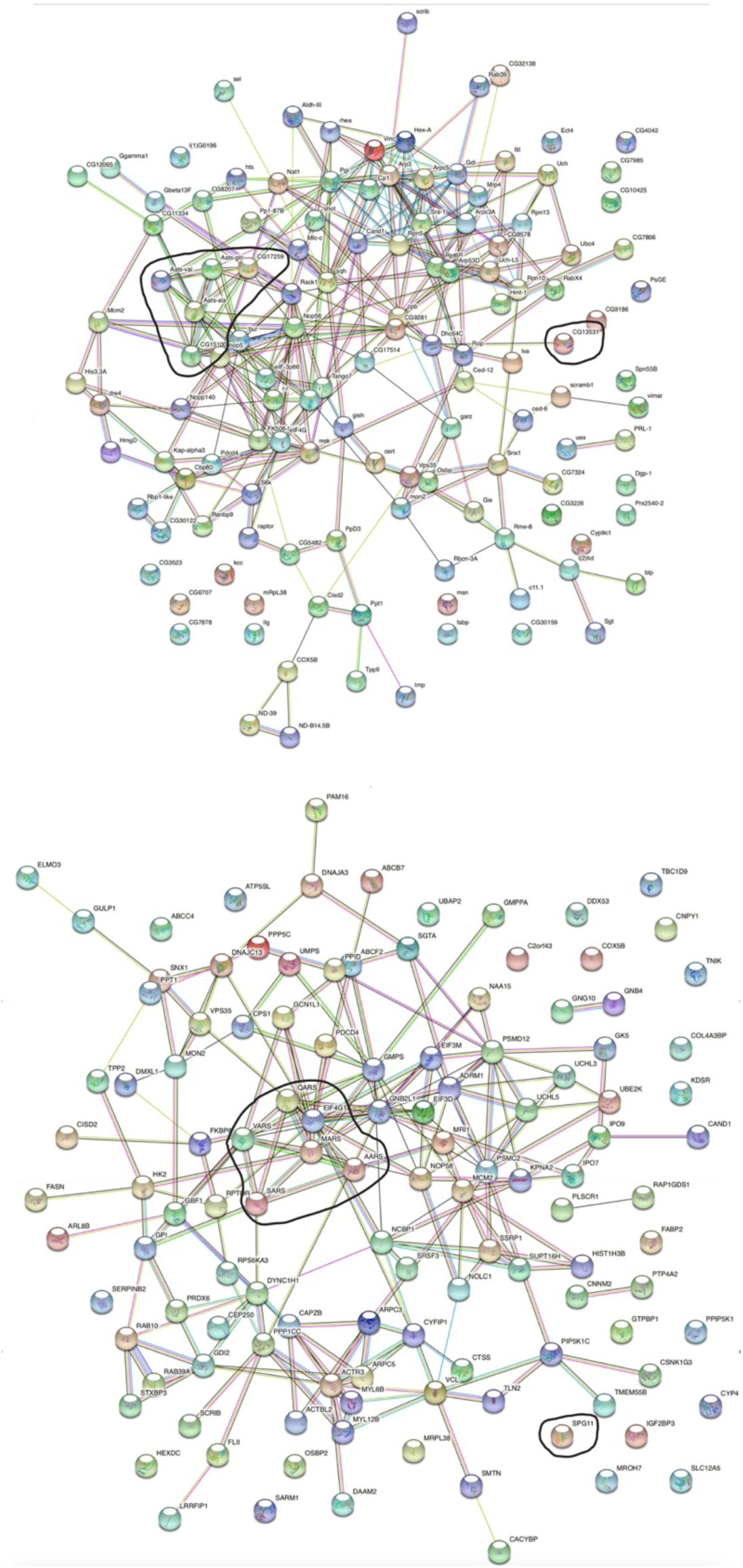
Interactions among Spg11_C binding partners found in both cell lines. Top: *Drosophila* Spg11_C partners. Bottom: human orthologs of *Drosophila* Spg11_C partners. Maps were generated using String (http://string-db.org). Aminoacyl-tRNA synthetases (including CG17259/SerRS and CG15100/MetRS) are grouped inside a freehand line; human EIF4G1 is an unrelated “bystander”. *Drosophila* and human CG13531/SPG11 are highlighted.

One unexpected class of SPG11-C binding partners were aminoacyl-tRNA synthetases, which charge specific tRNAs with the correct amino acid for translation. These were the highest ranking class of interactors with a specific enzymatic activity as a molecular function, and the highest ranking KEGG pathway (Table 4). Five of these were pulled down (Figure 6; Table 5). These included *Drosophila* homologs of methionyl-tRNA synthetase (MARS) and alanyl-tRNA synthetase (AARS), mutations in which are causative for Charcot-Marie-Tooth (CMT) Type 2 neuropathy (McLaughlin *et al*. 2012; Gonzalez *et al*. 2013; Timmerman *et al*. 2014; Wei *et al*. 2019), as well as serine-valine- and glutamine-tRNA synthetases (Table 5). The MARS ortholog was identified with a particularly high score in the inducible line (Table 1).

**Table 5.** Amino-acyl tRNA synthetase mass spectrometry results.

| PMT-GFP::Spg11-C |  |  |  |  |  |
| --- | --- | --- | --- | --- | --- |
| *Drosophila* Protein | Human Ortholog | CDS | Score | Matches | Sequences |
| MetRS | MARS | PA | 1147 | 26 | 14 |
| SerRS | SARS | PA | 341 | 6 | 4 |
| AlaRS | AARS | PB | 338 | 5 | 2 |
| ValRS | VARS | PB | 331 | 5 | 3 |
| GlnRS | QARS | PA | 125 | 4 | 3 |

Since the Vps16_C domain of Vps16 recruits the SM proteins Vps33a to the HOPS complex (Wartosch *et al*. 2015), we expected to find one or more SM proteins as binding partners of the spatacsin_C domain. *Drosophila* has 5 known SM proteins (Table 6). Three of these, Rop, Vps45 and Slh were identified in the output of the CuSO_4_-induced line (Figure 5B; Table 7). Rop was identified in both cell lines (Figure 5B; Table 3); we did not identify Vps33a (Carnation) or Vps33b as binding partners in either line.

**Table 6.** Members of the Sec1/Munc18 (SM) gene family in *Drosophila*.

| Gene Symbol | Gene Name | Also known as |
| --- | --- | --- |
| *car* | *carnation* | *Vps33, Vps33a, l(1)G0447* |
| *Rop* | *Ras opposite* | *l(3)64Ah, cs1, Sec1, Munc18* |
| *Slh* | *SLY-1 homologous* | *Sly1, d-sly* |
| *Vps33B* | *Vacuolar protein sorting 33B* |  |
| *Vps45* | *Vacuolar protein sorting 45* | *MEME(3R)-C* |

**Table 7.** Rop, Vps45, Slh and Spg15 mass spectrometry score results.

| PMT-GFP::Spg11-C |  |  |  |  |  |
| --- | --- | --- | --- | --- | --- |
| Gene Symbol | Protein | CDS | Score | Matches | Sequences |
| Rop | CG15811 | PA | 180 | 3 | 3 |
| Vps45 | CG8228 | PA | 122 | 2 | 2 |
| Slh | CG3539 | PC | 90 | 2 | 2 |

## Discussion

### Bioinformatic analyses predict functional roles of the N- and C-termini of SPG11

Our bioinformatic analyses make interesting predictions about two regions of the SPG11 protein family. First, our phylogenetic analyses show that the evolutionary presence of a conserved N-terminal region correlates strongly with the presence of the AP-5 adaptor complex, suggesting that binding to AP-5 is the only essential function of this N-terminal region. Second, the SPG11 C-terminus shows higher conservation than most other parts of SPG11, and both human and fly variants show predicted structural homology with the Vps16 C-terminus (Vps16_C, Pfam ID: PF04840) (Figure 2C). The model of homology is supported by many conserved residues in a multiple sequence alignment of spatacsin_C and Vps16_C domains (Figure 3A) and tertiary structure prediction by homology (Figure 3B).

The Vps16_C domain (Pfam ID: PF04840) mediates binding to Vps33 proteins (Baker *et al*. 2013; Graham *et al*. 2013). These belong to the Sec1/Munc18 (SM) protein family (Misura *et al*. 2000; Graham *et al*. 2013) and have three domains, arranged in an arch-like structure, with an internal cavity capable of binding to SNARE proteins (Misura *et al*. 2000; Baker *et al*. 2013; Toonen and Verhage 2003; Baker *et al*. 2015). SM proteins bind to SNARE proteins such as syntaxin and are important for membrane fusion in disparate pathways, including synapse neurotransmitter release (Wu *et al*. 1998; Harrison *et al*. 1994; Weimer and Richmond 2004), early endosomal tracking (Nielsen *et al*. 2000) (Morrison *et al*. 2008) and Golgi-ER tracking (Laufman *et al*. 2009). However, their precise role is not fully understood.

Humans and flies have two Vps16 proteins: Vps16a, which binds to Vps33a in the HOPS and CORVET complexes that promote fusion of endosomes or autophagosome to lysosomes (Bröcker *et al*. 2012; Urban *et al*. 2012; Solinger and Spang 2013), and Vps16b, which binds to Vps33b in the analogous but smaller CHEVI complex (Hunter *et al*. 2018; van der Beek *et al*. 2019). Therefore, the Spg11-C-terminal domain could potentially also bind to SM proteins; its divergence from known Vps16_C domains, and the relatively low conservation of Vps16 residues that bind to Vps33a, suggest that it is unlikely to bind to Vps33a or Vps33b, but it could potentially bind to other SM family proteins. The localization of the Spg11 C-terminus to acidic compartments (Figure 4A) could indicate a role for the Spg11-C-terminus in regulating some membrane fusion event(s) involving late endosomes, autophagosomes, autolysosomes, or lysosomes.

### SPG11 C-terminus interacts with SM proteins Rop, Slh and Vps45

Mass spectrometry analysis of the SPG11 C-terminus expressed in *Drosophila* Mel-2 cells was used to identify potential binding partners of the Spg11 C-terminus. This identified three of the five *Drosophila* SM proteins: Rop, Slh and Vps45, which were not found in the negative GFP control, tentatively indicating that the Spg11 C-terminus may possess SM protein binding ability.

Of the SM proteins identified, Rop was the highest scoring (Table 7). Rop is the homologue of yeast Sec1p (Wu *et al*. 1998), and is involved in synaptic vesicle exocytosis, synapse transmission and secretion (Wu *et al*. 1998; Harrison *et al*. 1994; Weimer and Richmond 2004); it does not appear to bind to Vps16a or Vps16b in *Drosophila* (Pulipparacharuvil *et al*. 2005). Vps45 is important in the endocytic pathway (Nielsen *et al*. 2000; Morrison *et al*. 2008). In both humans and flies, Vps45 is part of a complex with Rab5 and a FYVE-containing protein, Rabenosyn-5, to mediate vesicle fusion to form early endosomes (Nielsen *et al*. 2000; Morrison *et al*. 2008). In humans *VPS45* mutations are associated with congenital neutropenia, with human *VPS45* patients lacking lysosomes (Stepensky *et al*. 2013). Slh is the orthologue of yeast Sly1 (Toonen and Verhage 2003). In mammals, Sly1 is implicated in Golgi-to-ER retrograde transport (Laufman *et al*. 2009; Nogueira *et al*. 2014). At this stage we cannot tell whether the SM protein interactions of the Spatacsin_C domain are direct, or of functional biological significance. However, they do occur in a native *Drosophila* cellular environment, and they are therefore consistent with the model of a Vps16_C-like function for the Spatacsin_C domain.

### The Spg11 C-terminus interacts with aminoacyl tRNA synthetases

Mass spectrometry of GFP::Spg11-C pulled down five aminoacyl-tRNA (aa-tRNA) synthetases (Figure 6A), The aa-tRNA synthetases are essential for normal protein translation, by attaching the correct amino acid onto cognate tRNA molecules (Cusack 1997; Wallen and Antonellis 2013; Datta *et al*. 2009; Swairjo *et al*. 2004). However, mutations in at least two human aa-tRNA synthetases identified here, *MARS* and *AARS*, cause the axonal neuropathy, dominant Charcot-Marie-Tooth disease, type 2 (McLaughlin *et al*. 2012; Timmerman *et al*. 2014); furthermore, loss-of-function mutations in *SPG11* (Montecchiani *et al*. 2016); as well as mutations in the Spg11 binding partner, Rab7 (Gillingham *et al*. 2014), can cause CMT type 2 (Cogli *et al*. 2009). Therefore, mutations affecting SPG11, Rab7, and aa-tRNA synthetases might cause Charcot-Marie-Tooth disease by a common mechanism.

What kind of common mechanism could this be? CMT2 causality of aa-tRNA synthetases is a dominant trait that may be due to exposure of an internal surface by CMT2 mutations, and is independent of their enzymatic activity (Blocquel *et al*. 2017); the resulting dominant neurological CMT2 disease phenotype resembles an SPG11 recessive phenotype, suggesting that either SPG11 protein or aa-tRNA synthetases might antagonize each other. Aminoacyl tRNA synthetases have functions beyond translation, including amino acid sensing and autophagy regulation, by mediating mTOR activation (Han *et al*. 2012; Bonfils *et al*. 2012; Jewell *et al*. 2013). Further, when HeLa cells are treated with Leucine, LARS translocates to the lysosomal membrane where it acts as a GTPase-activating protein (GAP) for Rag GTPase (Han *et al*. 2012; Bonfils *et al*. 2012). Since SPG11 functions on autolysosomes to regenerate lysosomes, it could interact with aa-tRNA synthetases there. We therefore hypothesize that CMT2 pathology could be caused by defects in autophagosomal lysosome reformation as occurs in SPG11 loss-of-function (Varga *et al*. 2015), or by defects in lysosomal amino-acid-sensing machinery caused by gain-of-function mutations in aminoacyl-tRNA synthetases.

### Conclusions

Our analyses suggest that the sole essential function of the N-terminus of SPG11 is to bind the AP-5 adaptor complex, and support the model that its C-terminal spatacsin_C domain is a distantly related Vps_C domain, with roles in membrane trafficking. The recovery of several aminoacyl-tRNA synthetases as binding partners of this domain, suggests the possibility that mutations in human SPG11 or in aminoacyl-tRNA synthetases could give rise to Charcot-Marie-Tooth axonopathy by affecting a common cellular process in membrane trafficking.

## Acknowledgements

We thank Yuu Kimata and his group for help with *Drosophila* cell culture and establishment of cell lines, and the *Drosophila* Genomics Resource Center (supported by NIH Grant 2P40OD010949) for plasmids. This work was funded by a PhD studentship 861-792 from the UK Motor Neuron Disease Association and the Le Bouedec family to ALP, and by a grant from the Tom-Wahlig Stiftung to CJO’K.

