## Supplementary Text and Figures for "Distant homologies and domain conservation of the Hereditary Spastic Paraplegia protein SPG11/ALS5/spatacsin"

### Supplementary Information

| Accession Number | Description | Query Value | E Value | Identity |
|---|---|---|---|---|
| \|AIC61081.1\| | AP5B1 [synthetic construct] | 62% | 0 | 100% |
| \|AIC61080.1\| | AP5B1 [synthetic construct] | 25% | 1.00E-147 | 100% |
| \|XP_013380861.1\| | **PREDICTED: AP-5 complex subunit beta-1-like [Lingula anatina]** | 99% | 4.00E-47 | 25% |
| \|KXJ14650.1\| | AP-5 complex subunit beta-1 [Exaiptasia pallida] | 99% | 4.00E-36 | 24% |
| \|XP_009054849.1\| | **hypothetical protein LOTGIDRAFT_232378 [Lottia gigantea]** | 70% | 3.00E-21 | 22% |
| \|XP_011682012.1\| | PREDICTED: AP-5 complex subunit beta-1 [Strongylocentrotus purpuratus] | 43% | 2.00E-20 | 27% |
| \|XP_011428750.1\| | **PREDICTED: AP-5 complex subunit beta-1-like isoform X1 [Crassostrea gigas]** | 94% | 8.00E-20 | 23% |
| \|XP_011428751.1\| | **PREDICTED: AP-5 complex subunit beta-1-like isoform X2 [Crassostrea gigas]** | 82% | 9.00E-18 | 23% |
| \|XP_011682013.1\| | PREDICTED: AP-5 complex subunit beta-1-like [Strongylocentrotus purpuratus] | 21% | 8.00E-15 | 32% |
| \|XP_011406526.1\| | PREDICTED: AP-5 complex subunit beta-1-like [Amphimedon queenslandica] | 42% | 3.00E-09 | 26% |
| \|XP_003731064.1\| | PREDICTED: AP-5 complex subunit beta-1 [Strongylocentrotus purpuratus] | 17% | 2.00E-07 | 29% |
| \|KMZ62512.1\| | hypothetical protein ZOSMA_45G00540 [Zostera marina] | 32% | 2.00E-07 | 26% |
| \|XP_010654554.1\| | PREDICTED: uncharacterized protein LOC100249600 [Vitis vinifera] | 67% | 4.00E-07 | 24% |
| \|EFA83460.1\| | hypothetical protein PPL_03608 [Polysphondylium pallidum PN500] | 21% | 8.00E-07 | 23% |
| \|XP_013389819.1\| | **PREDICTED: AP-5 complex subunit beta-1-like [Lingula anatina]** | 18% | 8.00E-07 | 27% |
| \|XP_013777264.1\| | **PREDICTED: uncharacterized protein LOC106461943 [Limulus polyphemus]** | 17% | 9.00E-07 | 27% |
| \|KNA14629.1\| | hypothetical protein SOVF_105570 [Spinacia oleracea] | 26% | 4.00E-06 | 30% |
| \|KQL32241.1\| | hypothetical protein SETIT_0161441mg [Setaria italica] | 32% | 6.00E-06 | 29% |
| \|XP_004955298.1\| | PREDICTED: AP-5 complex subunit beta-1 [Setaria italica] | 32% | 6.00E-06 | 29% |
| \|XP_009418279.1\| | PREDICTED: uncharacterized protein LOC103998516 [Musa acuminata subsp. malaccensis] | 31% | 6.00E-06 | 32% |
| \|XP_006649198.1\| | PREDICTED: uncharacterized protein LOC102704955 [Oryza brachyantha] | 41% | 3.00E-05 | 24% |
| \|XP_012089641.1\| | PREDICTED: AP-5 complex subunit beta-1 [Jatropha curcas] | 30% | 5.00E-05 | 27% |
| \|GAU36940.1\| | hypothetical protein TSUD_62090 [Trifolium subterraneum] | 46% | 8.00E-05 | 22% |
| \|KXG31422.1\| | hypothetical protein SORBI_004G356000 [Sorghum bicolor] | 31% | 9.00E-05 | 29% |
| \|XP_002453090.1\| | hypothetical protein SORBIDRAFT_04g038250 [Sorghum bicolor] | 31% | 1.00E-04 | 30% |
| \|XP_015777333.1\| | PREDICTED: AP-5 complex subunit beta-1-like [Acropora digitifera] | 33% | 2.00E-04 | 27% |
| \|XP_013641080.1\| | PREDICTED: AP-5 complex subunit beta-1-like [Brassica napus] | 73% | 2.00E-04 | 23% |
| \|XP_010688752.1\| | PREDICTED: uncharacterized protein LOC104902615 [Beta vulgaris subsp. vulgaris] | 37% | 2.00E-04 | 26% |
| \|XP_004364858.1\| | **hypothetical protein CAOG_01990 [Capsaspora owczarzaki ATCC 30864]** | 16% | 2.00E-04 | 31% |
| \|XP_012756164.1\| | hypothetical protein SAMD00019534_042050 [Acytostelium subglobosum LB1] | 58% | 3.00E-04 | 20% |
| \|XP_007157305.1\| | hypothetical protein PHAVU_002G058700g [Phaseolus vulgaris] | 36% | 3.00E-04 | 25% |
| \|XP_006406446.1\| | hypothetical protein EUTSA_v10019942mg [Eutrema salsugineum] | 46% | 4.00E-04 | 24% |
| \|CEM21248.1\| | unnamed protein product [Vitrella brassicaformis CCMP3155] | 11% | 8.00E-04 | 33% |
| \|XP_012554216.1\| | **PREDICTED: AP-5 complex subunit beta-1-like [Hydra vulgaris]** | 34% | 8.00E-04 | 19% |

| Accession Number | Description | Query Value | E Value | Identity |
|---|---|---|---|---|
| XP_012554216.1 | **PREDICTED: AP-5 complex subunit beta-1-like [Hydra vulgaris]** | 100% | 0 | 100% |
| XP_002167159.1 | PREDICTED: thyrotroph embryonic factor-like isoform X1 [Hydra vulgaris] | 13% | 4.00E-77 | 97% |
| XP_012562208.1 | PREDICTED: thyrotroph embryonic factor-like isoform X2 [Hydra vulgaris] | 13% | 3.00E-76 | 97% |
| KXJ14650.1 | AP-5 complex subunit beta-1 [Exaiptasia pallida] | 45% | 9.00E-28 | 25% |
| XP_009054849.1 | **hypothetical protein LOTGIDRAFT_232378 [Lottia gigantea]** | 87% | 5.00E-27 | 24% |
| XP_001641289.1 | predicted protein [Nematostella vectensis] | 44% | 4.00E-24 | 25% |
| XP_013380861.1 | **PREDICTED: AP-5 complex subunit beta-1-like [Lingula anatina]** | 49% | 2.00E-22 | 26% |
| XP_014672098.1 | **PREDICTED: AP-5 complex subunit beta-1-like [Priapulus caudatus]** | 58% | 7.00E-22 | 23% |
| XP_011428751.1 | **PREDICTED: AP-5 complex subunit beta-1-like isoform X2 [Crassostrea gigas]** | 74% | 3.00E-18 | 22% |
| XP_011428750.1 | PREDICTED: AP-5 complex subunit beta-1-like isoform X1 [Crassostrea gigas] | 74% | 7.00E-18 | 22% |
| XP_011406526.1 | PREDICTED: AP-5 complex subunit beta-1-like [Amphimedon queenslandica] | 88% | 6.00E-16 | 22% |
| XP_011682013.1 | PREDICTED: AP-5 complex subunit beta-1-like [Strongylocentrotus purpuratus] | 25% | 1.00E-12 | 23% |
| XP_013068897.1 | **PREDICTED: AP-5 complex subunit beta-1-like [Biomphalaria glabrata]** | 86% | 2.00E-11 | 21% |
| XP_011682012.1 | PREDICTED: AP-5 complex subunit beta-1 [Strongylocentrotus purpuratus] | 17% | 1.00E-09 | 28% |
| XP_014780940.1 | **PREDICTED: AP-5 complex subunit beta-1-like [Octopus bimaculoides]** | 55% | 3.00E-09 | 24% |

| Accession Number | Description | Query Value | E Value | Identity |
|---|---|---|---|---|
| XP_013389819.1 | **PREDICTED: AP-5 complex subunit beta-1-like [Lingula anatina]** | 100% | 0 | 100% |
| XP_013380861.1 | PREDICTED: AP-5 complex subunit beta-1-like [Lingula anatina] | 92% | 1.00E-165 | 98% |
| XP_011682012.1 | PREDICTED: AP-5 complex subunit beta-1 [Strongylocentrotus purpuratus] | 92% | 2.00E-63 | 43% |
| ELT96156.1 | hypothetical protein CAPTEDRAFT_208254 [Capitella teleta] | 83% | 4.00E-40 | 37% |
| KXJ16464.1 | AP-5 complex subunit beta-1 [Exaiptasia pallida] | 86% | 1.00E-33 | 34% |
| KXJ14650.1 | AP-5 complex subunit beta-1 [Exaiptasia pallida] | 86% | 4.00E-31 | 34% |
| XP_011406526.1 | PREDICTED: AP-5 complex subunit beta-1-like [Amphimedon queenslandica] | 74% | 1.00E-29 | 35% |
| XP_004364858.1 | hypothetical protein CAOG_01990 [Capsaspora owczarzaki ATCC 30864] | 71% | 1.00E-28 | 36% |
| XP_015776887.1 | PREDICTED: AP-5 complex subunit beta-1-like [Acropora digitifera] | 79% | 1.00E-24 | 33% |
| XP_013780930.1 | PREDICTED: AP-5 complex subunit beta-1-like [Limulus polyphemus] | 69% | 6.00E-24 | 36% |
| XP_002114848.1 | **predicted protein [Trichoplax adhaerens]** | 54% | 1.00E-22 | 38% |
| XP_015908879.1 | **PREDICTED: AP-5 complex subunit beta-1-like [Parasteatoda tepidariorum]** | 68% | 3.00E-21 | 29% |

**Supplementary Figure 1. AP-5 beta-1 subunit BlastP searches.** Top Panel: BlastP searches (http://blast.ncbi.nlm.nih.gov) of reference sequences, excluding vertebrates, using AP-5 beta-1 subunit from (top list) humans (NCBI reference sequence: NP_612377.4), (middle list) *Hydra Vulgaris* (XP_012554216.1) and (lower list) *Lingula anatina* (XP_013389819.1), as queries to try and identify divergent orthologues across metazoans. Bold text indicates organisms present in SPG11 phylogenetic tree (Figure 1).

| Accession Number | Description | Query Value | E Value | Identity |
|---|---|---|---|---|
| AIC51767.1 | AP5S1 [synthetic construct] | 100% | 2.00E-141 | 100% |
| XP_003730604.1 | PREDICTED: AP-5 complex subunit sigma-1-like [Strongylocentrotus purpuratus] | 98% | 2.00E-36 | 40% |
| XP_013389295.1 | **PREDICTED: AP-5 complex subunit sigma-1-like [Lingula anatina]** | 98% | 5.00E-32 | 36% |
| XP_011438995.1 | **PREDICTED: AP-5 complex subunit sigma-1-like [Crassostrea gigas]** | 96% | 8.00E-32 | 38% |
| XP_002734372.1 | PREDICTED: AP-5 complex subunit sigma-1-like [Saccoglossus kowalevskii] | 96% | 1.00E-31 | 37% |
| XP_013418972.1 | **PREDICTED: AP-5 complex subunit sigma-1-like [Lingula anatina]** | 96% | 2.00E-31 | 37% |
| XP_015755081.1 | PREDICTED: AP-5 complex subunit sigma-1-like [Acropora digitifera] | 96% | 8.00E-31 | 38% |
| XP_009064272.1 | **hypothetical protein LOTGIDRAFT_196130 [Lottia gigantea]** | 96% | 5.00E-28 | 36% |
| XP_002587995.1 | hypothetical protein BRAFLDRAFT_125393 [Branchiostoma floridae] | 97% | 1.00E-26 | 37% |
| XP_002117045.1 | **hypothetical protein TRIADDRAFT_61078 [Trichoplax adhaerens]** | 77% | 4.00E-25 | 41% |
| XP_003384843.1 | PREDICTED: AP-5 complex subunit sigma-1-like [Amphimedon queenslandica] | 98% | 2.00E-20 | 31% |
| JAS63365.1 | hypothetical protein g.14383 [Cuerna arida] | 98% | 1.00E-13 | 30% |
| ELT87105.1 | **hypothetical protein CAPTEDRAFT_220165 [Capitella teleta]** | 98% | 2.00E-12 | 27% |
| JAS79307.1 | hypothetical protein g.17733 [Homalodisca liturata] | 98% | 3.00E-12 | 28% |
| JAS74495.1 | hypothetical protein g.17732 [Homalodisca liturata] | 98% | 8.00E-12 | 28% |
| XP_015910213.1 | **PREDICTED: AP-5 complex subunit sigma-1-like [Parasteatoda tepidariorum]** | 98% | 8.00E-11 | 27% |
| JAT17035.1 | hypothetical protein g.15854 [Graphocephala atropunctata] | 98% | 5.00E-10 | 26% |
| JAS16799.1 | hypothetical protein g.39886 [Clastoptera arizonana] | 96% | 4.00E-09 | 26% |
| XP_004342893.2 | hypothetical protein CAOG_07820 [Capsaspora owczarzaki ATCC 30864] | 48% | 5.00E-08 | 35% |
| KDR13731.1 | hypothetical protein L798_12129 [Zootermopsis nevadensis] | 49% | 8.00E-07 | 33% |
| ANQ07673.1 | Uncharacterized protein PCOAH_00022690 [Plasmodium coatneyi] | 49% | 2.00E-04 | 31% |
| SBT76639.1 | AP-5 complex subunit sigma-1, putative [Plasmodium falciparum] | 49% | 3.00E-04 | 28% |
| XP_013439434.1 | hypothetical protein, conserved [Eimeria necatrix] | 51% | 7.00E-04 | 36% |
| XP_013233372.1 | hypothetical protein, conserved [Eimeria tenella] | 44% | 0.001 | 39% |
| XP_002258714.1 | hypothetical protein, conserved in Plasmodium species [Plasmodium knowlesi strain H] | 98% | 0.001 | 24% |
| XP_012558475.1 | **PREDICTED: AP-5 complex subunit sigma-1-like [Hydra vulgaris]** | 97% | 0.001 | 23% |

| Accession Number | Description | Query Value | E Value | Identity |
|---|---|---|---|---|
| XP_009064272.1 | **hypothetical protein LOTGIDRAFT_196130 [Lottia gigantea]** | 100% | 1.00E-139 | 100% |
| XP_011438995.1 | **PREDICTED: AP-5 complex subunit sigma-1-like [Crassostrea gigas]** | 98% | 2.00E-68 | 51% |
| XP_013389295.1 | **PREDICTED: AP-5 complex subunit sigma-1-like [Lingula anatina]** | 97% | 9.00E-58 | 46% |
| XP_013418972.1 | **PREDICTED: AP-5 complex subunit sigma-1-like [Lingula anatina]** | 96% | 2.00E-56 | 46% |
| XP_002734372.1 | PREDICTED: AP-5 complex subunit sigma-1-like [Saccoglossus kowalevskii] | 96% | 4.00E-52 | 40% |
| XP_014768205.1 | **PREDICTED: uncharacterized protein LOC106867753 [Octopus bimaculoides]** | 98% | 5.00E-49 | 42% |
| XP_003730604.1 | PREDICTED: AP-5 complex subunit sigma-1-like [Strongylocentrotus purpuratus] | 96% | 3.00E-42 | 41% |
| XP_015755081.1 | PREDICTED: AP-5 complex subunit sigma-1-like [Acropora digitifera] | 99% | 4.00E-41 | 40% |
| XP_002587995.1 | hypothetical protein BRAFLDRAFT_125393 [Branchiostoma floridae] | 96% | 2.00E-34 | 36% |
| ELT87105.1 | **hypothetical protein CAPTEDRAFT_220165 [Capitella teleta]** | 98% | 2.00E-32 | 35% |
| XP_013094720.1 | **PREDICTED: uncharacterized protein LOC106078409 [Biomphalaria glabrata]** | 99% | 6.00E-28 | 36% |

**Supplementary Figure 2. AP-5 sigma-1 subunit BlastP searches.** Top Panel: BlastP searches (http://blast.ncbi.nlm.nih.gov) of reference sequences, excluding vertebrates, using AP-5 sigma-1 subunit from (top list) humans (NCBI reference sequence: NP_001191375.1), and (lower list) *Lottia gigantea* (NCBI reference sequence: XP_00906064272.1) as queries to try and identify divergent orthologues across metazoans. Bold text indicates organisms present in SPG11 phylogenetic tree (Figure 1).

| Accession Number | Description | Query Value | E Value | Identity |
|---|---|---|---|---|
| ADZ15988.1 | KIAA0415 [synthetic construct] | 55% | 0 | 100% |
| XP_013380903.1 | **PREDICTED: AP-5 complex subunit zeta-1-like [Lingula anatina]** | 99% | 8.00E-123 | 31% |
| XP_013416069.1 | **PREDICTED: AP-5 complex subunit zeta-1-like [Lingula anatina]** | 99% | 2.00E-122 | 31% |
| XP_011438415.1 | **PREDICTED: AP-5 complex subunit zeta-1-like [Crassostrea gigas]** | 98% | 3.00E-120 | 32% |
| EKC38143.1 | **hypothetical protein CGI_10019308 [Crassostrea gigas]** | 96% | 4.00E-118 | 32% |
| XP_795875.1 | PREDICTED: AP-5 complex subunit zeta-1 [Strongylocentrotus purpuratus] | 94% | 2.00E-117 | 33% |
| XP_003382874.1 | PREDICTED: AP-5 complex subunit zeta-1-like [Amphimedon queenslandica] | 96% | 1.00E-115 | 34% |
| XP_796686.4 | PREDICTED: AP-5 complex subunit zeta-1 [Strongylocentrotus purpuratus] | 94% | 4.00E-107 | 32% |
| XP_015771760.1 | PREDICTED: AP-5 complex subunit zeta-1-like [Acropora digitifera] | 94% | 8.00E-95 | 38% |
| XP_014770406.1 | **PREDICTED: AP-5 complex subunit zeta-1-like [Octopus bimaculoides]** | 98% | 2.00E-94 | 28% |
| XP_013067408.1 | **PREDICTED: AP-5 complex subunit zeta-1-like [Biomphalaria glabrata]** | 98% | 4.00E-91 | 29% |
| XP_002595760.1 | hypothetical protein BRAFLDRAFT_117560 [Branchiostoma floridae] | 70% | 7.00E-91 | 34% |
| XP_013782855.1 | **PREDICTED: AP-5 complex subunit zeta-1-like [Limulus polyphemus]** | 83% | 4.00E-86 | 31% |
| KOF91423.1 | **hypothetical protein OCBIM_22008898mg [Octopus bimaculoides]** | 79% | 3.00E-83 | 29% |
| KFM68184.1 | hypothetical protein X975_21262 [Stegodyphus mimosarum] | 61% | 5.00E-79 | 32% |
| ELU06156.1 | **hypothetical protein CAPTEDRAFT_227527 [Capitella teleta]** | 76% | 1.00E-73 | 33% |
| KXJ13612.1 | AP-5 complex subunit zeta-1 [Exaiptasia pallida] | 53% | 2.00E-72 | 36% |
| XP_006817102.1 | PREDICTED: AP-5 complex subunit zeta-1-like [Saccoglossus kowalevskii] | 45% | 4.00E-72 | 37% |
| XP_009045402.1 | **hypothetical protein LOTGIDRAFT_230221 [Lottia gigantea]** | 98% | 1.00E-68 | 34% |
| XP_002109736.1 | **hypothetical protein TRIADDRAFT_52911 [Trichoplax adhaerens]** | 90% | 6.00E-68 | 27% |
| KDR14925.1 | hypothetical protein L798_11231 [Zootermopsis nevadensis] | 77% | 3.00E-65 | 29% |
| XP_001639990.1 | predicted protein [Nematostella vectensis] | 93% | 3.00E-63 | 31% |
| XP_015903436.1 | **PREDICTED: AP-5 complex subunit zeta-1-like [Parasteatoda tepidariorum]** | 60% | 4.00E-55 | 30% |

| Accession Number | Description | Query Value | E Value | Identity |
|---|---|---|---|---|
| XP_015903436.1 | **PREDICTED: AP-5 complex subunit zeta-1-like [Parasteatoda tepidariorum]** | 100% | 0 | 100% |
| KFM68184.1 | hypothetical protein X975_21262 [Stegodyphus mimosarum] | 55% | 2.00E-154 | 66% |
| XP_013782855.1 | **PREDICTED: AP-5 complex subunit zeta-1-like [Limulus polyphemus]** | 94% | 6.00E-109 | 37% |
| XP_014770406.1 | **PREDICTED: AP-5 complex subunit zeta-1-like [Octopus bimaculoides]** | 85% | 2.00E-69 | 30% |
| XP_796686.4 | PREDICTED: AP-5 complex subunit zeta-1 [Strongylocentrotus purpuratus] | 87% | 9.00E-69 | 31% |
| XP_795875.1 | PREDICTED: AP-5 complex subunit zeta-1 [Strongylocentrotus purpuratus] | 87% | 8.00E-66 | 30% |
| KOF91423.1 | **hypothetical protein OCBIM_22008898mg [Octopus bimaculoides]** | 80% | 3.00E-65 | 29% |
| XP_011438415.1 | **PREDICTED: AP-5 complex subunit zeta-1-like [Crassostrea gigas]** | 82% | 8.00E-65 | 31% |
| EKC38143.1 | **hypothetical protein CGI_10019308 [Crassostrea gigas]** | 82% | 8.00E-65 | 31% |
| ADZ15988.1 | KIAA0415 [synthetic construct] | 74% | 1.00E-55 | 31% |

**Supplementary Figure 3. AP-5 zeta-1 subunit BlastP searches.** Top Panel: BlastP searches (http://blast.ncbi.nlm.nih.gov) of reference sequences, excluding vertebrates, using AP-5 zeta-1 subunit from (top list) humans (NCBI reference sequence: NP_055670.1), and (lower list) *Parasteatoda tepidariorum* (NCBI reference sequence: XP_015903436.1) as queries to try and identify divergent orthologues across metazoans. Bold text indicates organisms present in SPG11 phylogenetic tree (Figure 1).

| Accession Number | Description | Query Value | E Value | Identity |
|---|---|---|---|---|
| AIC60393.1 | MUDENG [synthetic construct] | 48% | 2.00E-173 | 100% |
| XP_002604479.1 | hypothetical protein BRAFLDRAFT_79220 [Branchiostoma floridae] | 98% | 1.00E-120 | 42% |
| XP_014663260.1 | **PREDICTED: AP-5 complex subunit mu-1-like [Priapulus caudatus]** | 93% | 3.00E-108 | 38% |
| XP_011446776.1 | PREDICTED: AP-5 complex subunit mu-1-like [Crassostrea gigas] | 97% | 9.00E-103 | 37% |
| XP_002731573.1 | PREDICTED: AP-5 complex subunit mu-1-like [Saccoglossus kowalevskii] | 97% | 5.00E-102 | 37% |
| XP_013392723.1 | **PREDICTED: AP-5 complex subunit mu-1-like [Lingula anatina]** | 99% | 2.00E-95 | 38% |
| XP_013399534.1 | **PREDICTED: AP-5 complex subunit mu-1-like [Lingula anatina]** | 99% | 1.00E-94 | 37% |
| XP_001632221.1 | predicted protein [Nematostella vectensis] | 93% | 1.00E-90 | 38% |
| KXJ19112.1 | AP-5 complex subunit mu-1 [Exaiptasia pallida] | 99% | 1.00E-85 | 35% |
| XP_013784486.1 | **PREDICTED: AP-5 complex subunit mu-1-like [Limulus polyphemus]** | 98% | 2.00E-80 | 34% |
| XP_014778639.1 | **PREDICTED: AP-5 complex subunit mu-1-like [Octopus bimaculoides]** | 97% | 3.00E-79 | 33% |
| XP_004349404.2 | hypothetical protein CAOG_02654 [Capsaspora owczarzaki ATCC 30864] | 98% | 2.00E-78 | 32% |
| KJE91526.1 | hypothetical protein CAOG_002654 [Capsaspora owczarzaki ATCC 30864] | 98% | 1.00E-77 | 32% |
| XP_015770274.1 | PREDICTED: AP-5 complex subunit mu-1-like [Acropora digitifera] | 86% | 2.00E-76 | 35% |
| XP_782538.2 | PREDICTED: AP-5 complex subunit mu-1 [Strongylocentrotus purpuratus] | 64% | 6.00E-69 | 39% |
| XP_002415549.1 | conserved hypothetical protein [Ixodes scapularis] | 84% | 1.00E-64 | 34% |
| XP_015927347.1 | **PREDICTED: AP-5 complex subunit mu-1-like [Parasteatoda tepidariorum]** | 98% | 4.00E-64 | 31% |
| EKC24584.1 | Kinesin-like protein KIF6 [Crassostrea gigas] | 64% | 4.00E-63 | 39% |
| XP_002117995.1 | **hypothetical protein TRIADDRAFT_62031 [Trichoplax adhaerens]** | 93% | 3.00E-62 | 30% |
| XP_004353035.1 | adaptor complexes medium subunit family protein [Acanthamoeba castellanii str. Neff] | 57% | 5.00E-61 | 39% |
| XP_013064051.1 | **PREDICTED: AP-5 complex subunit mu-1-like [Biomphalaria glabrata]** | 73% | 5.00E-61 | 36% |
| XP_012553969.1 | **PREDICTED: AP-5 complex subunit mu-1-like isoform X1 [Hydra vulgaris]** | 97% | 5.00E-59 | 31% |

| Accession Number | Description | Query Value | E Value | Identity |
|---|---|---|---|---|
| XP_015927347.1 | **PREDICTED: AP-5 complex subunit mu-1-like [Parasteatoda tepidariorum]** | 100% | 0 | 100% |
| KFM57535.1 | MHD domain-containing death-inducing protein [Stegodyphus mimosarum] | 63% | 3.00E-118 | 59% |
| XP_013784486.1 | **PREDICTED: AP-5 complex subunit mu-1-like [Limulus polyphemus]** | 97% | 2.00E-98 | 38% |
| XP_011446776.1 | PREDICTED: AP-5 complex subunit mu-1-like [Crassostrea gigas] | 96% | 2.00E-80 | 32% |
| XP_002731573.1 | PREDICTED: AP-5 complex subunit mu-1-like [Saccoglossus kowalevskii] | 96% | 2.00E-78 | 34% |
| XP_002604479.1 | hypothetical protein BRAFLDRAFT_79220 [Branchiostoma floridae] | 96% | 9.00E-73 | 34% |
| XP_013399534.1 | **PREDICTED: AP-5 complex subunit mu-1-like [Lingula anatina]** | 96% | 1.00E-68 | 32% |
| XP_013392723.1 | **PREDICTED: AP-5 complex subunit mu-1-like [Lingula anatina]** | 96% | 4.00E-68 | 32% |
| XP_014663260.1 | **PREDICTED: AP-5 complex subunit mu-1-like [Priapulus caudatus]** | 91% | 1.00E-65 | 31% |
| XP_002415549.1 | conserved hypothetical protein [Ixodes scapularis] | 87% | 4.00E-63 | 31% |
| XP_014778639.1 | **PREDICTED: AP-5 complex subunit mu-1-like [Octopus bimaculoides]** | 96% | 1.00E-61 | 30% |
| KXJ19112.1 | AP-5 complex subunit mu-1 [Exaiptasia pallida] | 96% | 1.00E-56 | 30% |
| XP_001632221.1 | predicted protein [Nematostella vectensis] | 91% | 1.00E-52 | 30% |
| XP_015770274.1 | PREDICTED: AP-5 complex subunit mu-1-like [Acropora digitifera] | 82% | 3.00E-50 | 30% |
| KJE91526.1 | hypothetical protein CAOG_002654 [Capsaspora owczarzaki ATCC 30864] | 77% | 1.00E-49 | 29% |
| XP_004349404.2 | hypothetical protein CAOG_02654 [Capsaspora owczarzaki ATCC 30864] | 77% | 2.00E-49 | 29% |
| EKC24584.1 | Kinesin-like protein KIF6 [Crassostrea gigas] | 62% | 2.00E-46 | 37% |
| XP_011408564.1 | PREDICTED: AP-5 complex subunit mu-1-like [Amphimedon queenslandica] | 62% | 1.00E-44 | 35% |
| XP_012553969.1 | **PREDICTED: AP-5 complex subunit mu-1-like isoform X1 [Hydra vulgaris]** | 96% | 3.00E-43 | 27% |

**Supplementary Figure 4. AP-5 mu-1 subunit BlastP searches.** Top Panel: BlastP searches (http://blast.ncbi.nlm.nih.gov) of reference sequences, excluding vertebrates, using AP-5 sigma-1 subunit from (top list) humans (NCBI reference sequence: NP_060699.3), and (lower list) *Parasteatoda tepidariorum* (NCBI reference sequence: XP_015927347.1) as queries to try and identify divergent orthologues across metazoans. Bold text indicates organisms present in SPG11 phylogenetic tree (Figure 1).

**Supplementary Figure 5. HH-pred output of a sequence search using an alignment of Spg11_C domains as a query.**

Alignments of a diverse group of Spg11 C-terminal domains, from *Apis*, *Danio*, *Drosophila*, *Gallus*, *Homo*, *Trichoplax* were generated using T-Coffee (Notredame *et al.* 2000). This alignment was used as a query in a HHpred search, and the highest 25 hits are listed. Alignments are presented between *Apis* Spg11 C-terminus and selected members of the Spg11, Vps16a and Vps16b families.

### HHpred

| Nr | Hit | Name | Probability | E-value | SS | Cols | Target Length |
|---|---|---|---|---|---|---|---|
| 1 | XP_016878125.1 | PREDICTED: spatacsin isoform X5 [Homo sapiens] | 100 | 5.8e-86 | 34.2 | 296 | 1402 |
| 2 | XP_006720763.1 | PREDICTED: spatacsin isoform X2 [Homo sapiens] | 100 | 8.5e-84 | 33.5 | 296 | 2395 |
| 3 | NP_079413.3 | spatacsin isoform 1 [Homo sapiens] | 100 | 2.3e-83 | 34 | 296 | 2443 |
| 4 | NP_001286748.1 | uncharacterized protein Dmel_CG13531, isoform C [Drosophila melanogaster] | 100 | 2.5e-83 | 32.5 | 306 | 1815 |
| 5 | NP_611726.1 | uncharacterized protein Dmel_CG13531, isoform B [Drosophila melanogaster] | 100 | 2.5e-83 | 32.5 | 306 | 1815 |
| 6 | NP_001153699.1 | spatacsin isoform 2 [Homo sapiens] | 100 | 3e-82 | 33.9 | 296 | 2330 |
| 7 | XP_016878123.1 | PREDICTED: spatacsin isoform X1 [Homo sapiens] | 100 | 2.2e-66 | 28.8 | 260 | 2407 |
| 8 | XP_011535368.1 | PREDICTED: spermatogenesis-defective protein 39 homolog isoform X1 [Homo sapiens] | 89.51 | 73 | 18.4 | 227 | 462 |
| 9 | XP_016877068.1 | PREDICTED: spermatogenesis-defective protein 39 homolog isoform X1 [Homo sapiens] | 89.51 | 73 | 18.4 | 227 | 462 |
| 10 | NP_649877.1 | vacuolar protein sorting 16A [Drosophila melanogaster] | 86.55 | 140 | 22.4 | 252 | 833 |
| 11 | NP_651731.1 | vacuolar protein sorting 16B [Drosophila melanogaster] | 81.55 | 180 | 20.2 | 242 | 447 |
| 12 | NP_001262478.1 | elongator complex protein 1, isoform C [Drosophila melanogaster] | 74.68 | 230 | 13.6 | 227 | 1256 |
| 13 | NP_650098.1 | elongator complex protein 1, isoform A [Drosophila melanogaster] | 69.55 | 340 | 13.5 | 227 | 1252 |
| 14 | XP_016875043.1 | PREDICTED: DCC-interacting protein 13-beta isoform X6 [Homo sapiens] | 69.52 | 350 | 12.8 | 126 | 617 |
| 15 | NP_001262477.1 | elongator complex protein 1, isoform B [Drosophila melanogaster] | 68.01 | 310 | 12.9 | 192 | 1253 |
| 16 | NP_001006635.1 | rho GTPase-activating protein 17 isoform 1 [Homo sapiens] | 62.35 | 690 | 18.1 | 143 | 881 |
| 17 | NP_001180243.1 | spermatogenesis-defective protein 39 homolog isoform 1 [Homo sapiens] | 60.59 | 590 | 21 | 250 | 493 |
| 18 | NP_001180244.1 | spermatogenesis-defective protein 39 homolog isoform 1 [Homo sapiens] | 60.59 | 590 | 21 | 250 | 493 |
| 19 | NP_001180246.1 | spermatogenesis-defective protein 39 homolog isoform 1 [Homo sapiens] | 60.59 | 590 | 21 | 250 | 493 |
| 20 | NP_071350.2 | spermatogenesis-defective protein 39 homolog isoform 1 [Homo sapiens] | 60.59 | 590 | 21 | 250 | 493 |
| 21 | XP_016876615.1 | PREDICTED: zinc finger FYVE domain-containing protein 26 isoform X3 [Homo sapiens] | 57.43 | 1200 | 24.3 | 281 | 2042 |
| 22 | XP_016868887.1 | PREDICTED: eukaryotic translation initiation factor 3 subunit E isoform X1 [Homo sapiens] | 51.84 | 430 | 9.8 | 114 | 480 |
| 23 | NP_610064.2 | outer segment 5 [Drosophila melanogaster] | 50.37 | 450 | 9.2 | 133 | 775 |
| 24 | NP_001180245.1 | spermatogenesis-defective protein 39 homolog isoform 2 [Homo sapiens] | 50.31 | 810 | 20.5 | 252 | 444 |
| 25 | NP_072097.2 | vacuolar protein sorting-associated protein 16 homolog isoform 1 [Homo sapiens] | 50.11 | 1000 | 24.9 | 276 | 839 |

### *Alignments*

1. XP_016878125.1 **PREDICTED: spatacsin isoform X5 [Homo sapiens]**

Probability: 100%, E-value: 5.8e-86, Score: 699.89, Aligned cols: 296, Identities: 30%, Similarity: 0.501,

```
Q ss_pred               CcchhHHHHHHHHHHHHhccchHhHHHHHHHHHhchHHHHHHccCCCccccCHHHHHHHHHHHCCCChHHHHHHHHHHHH
Q Apis_Spg11       1    NILQNLKYWTLLVRLVTGVGRFTEMNYIFQILKENDQFEFLLGKGLNKVTGLETALLDFLKHHCPENKELFTLVALHFRL   80 (319)
Q Consensus        1    n~L~~~~~~~~l~VRLltGIgRy~EM~YiFdiL~en~qFE~Ll~k~~~~~~~~L~tAlLdylKr~~P~d~Ek~~mvAL~FsM   80 (319)
                        ++|++.+.|+|||||||||||+||+||||+|++|||||.||+|+.+++++||+||+||||||+|+|+|+|+|+||||+|
T Consensus     1107 ~~l~~~~~~~~llvRLltGigry~em~yifd~L~~n~qfE~LL~k~~d~~~~lk~AlldyLk~~~P~d~e~~~mval~F~M   1186 (1402)
T XP_016878125.1 1107 NHLAPSEEYGLVVRLLTGIGRYNEMTYIFDLLHKKHYFEVLMRKKLDPSGTLKTALLDYIKRCRPGDSEKHNMIALCFSM   1186 (1402)
T ss_pred               cccCChhhhHhhhhHHhcCCCHHHHHHHHHHHHHcccHHHHhhcccCCCchHHHHHHHHHHHhCCCCHHHHHHHHHHHHh

Q ss_pred               HHHHHHHhHHHHHHHHHHHHhhhhHHHHhhhcchHhhHhhcchHHHHHHHHHHHHHHHHHHHHHhHHHHHHHHHHHHHHH
Q Apis_Spg11       81   YHELALMWENEAKDLIKTIISNATKEYNKLQSNVQHEIKFIKTEYIQKQLQLIITNYTHATEYYLQANKLNLASQCSDQV   160 (319)
Q Consensus        81   ~rEige~~e~~A~~~lk~I~sq~~~~~~~~Lq~~~~~~~~~~~~~~~~~~~kk~L~~~~t~~tdAaE~Y~kd~c~r~A~qc~~~a   160 (319)
                        |||||+|||++|+++++.|++++|++ ++++ ++++++.|++|+++|+||||+|++|+|+++|++|.++|
T Consensus     1187 ~~Eia~~~e~~A~~~l~~i~~~~~~~~--~~~~----------~~~~~~~~L~~am~~~~~Aae~y~~~~~~~~~~~A~~C~~~a   1254 (1402)
T XP_016878125.1 1187 CREIGENHEAAARIQLKLIESQPWED--SLKD----------GHQLKQLLLKALTLMLDAAESYAKDSCVRQAQHCQRLT   1254 (1402)
T ss_pred               HHHHHHHHHHHHHHHHHHHHcCcchh--hcCC----------hHHHHHHHHHHHHHHHHHHHHHhhhHHHHHHHHHHHHH

Q ss_pred               HHHHHHHHhhccCCCCcceEEeecCCHHHHHHHHHcCCchHHHHHHHHHhccCCcHHHHHHHHHHHhCCchHHHHHHHhc
Q Apis_Spg11       161  QLVALQLSLFNTTSYNQQVICILNLKPEDIDKVLCHNLSFSQCFIIVHAYNHHVDWANLIYNHCILNGETKYLKDFITVN   240 (319)
Q Consensus        161  ~Lv~lQl~lln~~~~~~~~~~~~~~~~~~Inl~~~~l~~~~~~~~~~~~~f~Qa~iva~AY~~~~dWae~iy~qvI~~g~f~YL~eF~~~~   240 (319)
                        +||+|||+++|++. .+++|||++++++++++++.+|+|+||+|||+|||+++||+++||+|||.+||++|++||++++
T Consensus     1255 ~LvaLQi~~~~~~~~----~~~iinL~~~~~~~~~~i~~~~~f~ealiva~aY~~~~~W~~~l~~q~v~~~~~~Yl~ef~~~~   1330 (1402)
T XP_016878125.1 1255 KLITLQIHFLNTGQ----NTMLINLGRHKLMDCILALPRFYQASIVAEAYDFVPDWAEILYQQVILKGDFNYLEEFKQQR   1330 (1402)
T ss_pred               HHHHHHHHcccCCC----CceEEeCCHHHHHHHHHhCCChHHHHHHHHHHcCCCchHHHHHHHHHHHhCChHHHHHHHhcC

Q ss_pred               CCCHHHHHHHHHHHhccCCcchHHHHHHHHHHHHHHhchHHHHHHHHhCcHHHHHHHhhCcccchhhhhhhhh
Q Apis_Spg11       241  KLTPSLVEDCVRRYRLEKSITHTMTNNMKILISELSDVECKYVLASQLGFKNIVEAMLNNPMIGAYLKDTVWK   313 (319)
Q Consensus        241  ~l~~s~~edi~kky~~~kp~~~~~~~~~~nmk~Li~~~~dV~~~Y~LA~~lgF~~IV~~LLnd~~~g~~LkDt~~~   313 (319)
                        +++|++++||+|+|++++|+ +.+.+|||+|+++|+||+++|+||+++||+|||++|++||++|+|||||+++
T Consensus     1331 ~l~~~~~~d~~~~~~~~~~~~~-~~~~~~~~~~ll~~~~d~~~~y~la~~~gf~d~~~~l~~~~~~~~~~~l~d~~~~   1402 (1402)
T XP_016878125.1 1331 LLKSSIFEEISKKYKQHQPT-DMVMENLKKLLTYCEDVYLYYKLAYEHKFYEIVNVLLKDPQTGCCLKDMLAG   1402 (1402)
T ss_pred               CCChhHHHHHHHHHHhcCCc-HHHHHHHHHHHHHCChHHHHHHHHHHcCCHHHHHHHhhCcccccchhHhhcC
```

---

4. NP_001286748.1 **uncharacterized protein Dmel_CG13531, isoform C [Drosophila melanogaster]**

Probability: 100%, E-value: 2.5e-83, Score: 694.07, Aligned cols: 306, Identities: 35%, Similarity: 0.581,

```
Q ss_pred               CcchhHHHHHHHHHHHHhccchHhHHHHHHHHHhchHHHHHHccCC-CcccCHHHHHHHHHHHHCCCC-hHHHHHHHHHH
Q Apis_Spg11       1    NILQNLKYWTLLVRLVTGVGRFTEMNYIFQILKENDQFEFLLGKGL-NKVTGLETALLDFLKHHCPEN-KELFTLVALHF   78 (319)
Q Consensus        1    n~L~~~~~~~~l~VRLltGIgRy~EM~YiFdiL~en~qFE~Ll~k~~-~~~~~~L~tAlLdylKr~~P~d-~Ek~~mvAL~F   78 (319)
                        ++|++.+.|+|||||||||||+||+||||+|++|||||.||+|+. +++.+||+||+||||||+|+| +|+++|+||||
T Consensus     1492 ~~l~~~~~~~~llVRLltGigry~em~yifd~L~~n~qfE~LL~k~~~~~~~~~~~Lk~AlldyLk~~~P~d~~e~~~mval~F   1571 (1815)
T NP_001286748.1 1492 TKLAQAKSWSLIVRLLIGIARYREMFYCFDSLIENEQFESLLGQFDEDQKGGLRQAILSYLREYQPKNGKELLRLAALHF   1571 (1815)
T ss_pred               HHHhhccccHHHHHHHHccCcHHHHHHHHHHHHHcchHHHHHhcCcccccchHHHHHHHHHHHHHCCCCHHHHHHHHHHHH

Q ss_pred               HHHHHHHHhHHHHHHHHHHHHHHh-hhhHHHHhhcchHhhHhhcchHHHHHHHHHHHHHHHHHHHHHhHHHHHHHHHHHH
Q Apis_Spg11       79   RLYHELALMWENEAKDLIKTIISN-ATKEYNKLQSNVQHEIKFIKTEYIQKQLQLIITNYTHATEYYLQANKLNLASQCS   157 (319)
Q Consensus        79   sM~rEige~~e~~A~~~lk~I~sq-~~~~~~~~~Lq~~~~~~~~~~~~~~~~~~~~kk~L~~~~t~~tdAaE~Y~kd~c~r~A~qc~   157 (319)
                        +||||||+|||++|+++++.|.++ +|++ ++++ ++++++.|++|+++|+||||+|+++||+++|++|.
T Consensus     1572 ~M~~Eia~~~e~~A~~~l~~~~~~~~~~~~~~--~~~~----------~~~~~~~~L~~am~~~~~Aae~y~~~~~~~~~~~A~~c~   1639 (1815)
T NP_001286748.1 1572 LMYKELAEMWTTEAQEIVTKIQAFAAQSN--KLKC----------SVEVQTLLQQALENYTHATENYLLDNKLLLAQQSV   1639 (1815)
T ss_pred               HhHHHHHHHHHHHHHHHHHHHHHHHHhcc--ccCC----------CHHHHHHHHHHHHHHHHHHHHHHHhcchHHHHHHHH

Q ss_pred               HHHHHHHHHHHhhccCCCC-------cceEEeecC-CHHHHHHHHHcCCchHHHHHHHHHhccCCcHHHHHHHHHHHhCC
Q Apis_Spg11       158  DQVQLVALQLSLFNTTSYN-------QQVICILNL-KPEDIDKVLCHNLSFSQCFIIVHAYNHHVDWANLIYNHCILNGE   229 (319)
Q Consensus        158  ~~a~Lv~lQl~lln~~~~~~-------~~~~~~~Inl-~~~~l~~~~~~~~~~~~~f~Qa~iva~AY~~~~dWae~iy~qvI~~g~   229 (319)
                        ++|+||+|||+++|.+..+ ++.+++||| ++++++++++++.+|+|+||+|||+|||+++||+++||+|||.+|+
T Consensus     1640 ~~a~LvaLQi~~~~~~~~~~~~~~~~~~~~~~~~~~~~~~~vlnL~s~~~~~~~~~i~~~~~f~qalIva~AY~~~~~W~~~l~~q~v~~~~   1719 (1815)
T NP_001286748.1 1640 SRAELMAMQLDLCNKALEKRHSNNANHLCVSVIGVRSREQFRELVNVHLSVPQALILSRAYGYDISWSEAILSQFVVLQG   1719 (1815)
T ss_pred               HHHHHHHHHHHhcHHHHHhcccCCccceeEEEccCCCHHHHHHHHHHcccchHHHHHHHHHHCCCCchHHHHHHHHhhcCC

Q ss_pred               chHHHHHHHhcCCCHHHHHHHHHHHhcc---CCcchHHHHHHHHHHHHHHhchHHHHHHHHhCcHHHHHHHhhCccccchh
Q Apis_Spg11       230  TKYLKDFITVNKLTPSLVEDCVRRYRLE---KSITHTMTNNMKILISELSDVECKYVLASQLGFKNIVEAMLNNPMIGAY   306 (319)
Q Consensus        230  f~YL~eF~~~~~l~~s~~edi~kky~~~---kp~~~~~~~~~~nmk~Li~~~~dV~~~Y~LA~~lgF~~IV~~LLnd~~~g~~   306 (319)
                        ++||+||+++++++|++++||+|+|+|+.+ ++++..|.+|||+|+++|+||+++|+||+++||+|||++|++| ++|+|
T Consensus     1720 ~~YL~ef~~~~~l~~~~~~~~~i~k~y~~~~~~~~~~~~~~~~~~~~~~~~~~ll~~~~d~~~~y~LA~~~gf~d~v~~ll~~-~~~~y   1798 (1815)
T NP_001286748.1 1720 VNYLQEYLCHQRINDDVIEQIVKGYLLHIQSNATNSKQEESMVQLVGLIKSVVLKYKLASILGFKSIVMSLIND-SSVYY   1798 (1815)
T ss_pred               hHHHHHHHhcCCCCHHHHHHHHHHHHHHHhcCCCCCChhHHHHHHHHHHCccHHHHHHHHHHhCCHHHHHHHHhcC-Cccee

Q ss_pred               hhhhhhhcccCCC
Q Apis_Spg11       307  LKDTVWKKGYNAT   319 (319)
Q Consensus        307  LkDt~~~~~~~~~~~t   319 (319)
                        ||||+|||+.+++
T Consensus     1799 l~D~~~~~~~~~~~~~   1811 (1815)
T NP_001286748.1 1799 LRDTNFGRTDFHT   1811 (1815)
T ss_pred               cchhccccccccC
```

8. XP_011535368.1 PREDICTED: spermatogenesis-defective protein 39 homolog isoform X1 [Homo sapiens]
Probability: 89.51%, E-value: 73, Score: 35.22, Aligned cols: 227, Identities: 14%, Similarity: 0.112,

```
Q ss_pred               HHHHHHhccchHhHHHHHHHHHHhchHHHHHHccCCCcccCHHHHHHHHHHHHCCCChHHHHHHHHHHHHHHHHHHHhHHHH
Q Apis_Spg11       12  LVRLVTGVGRFTEMNYIFQILKENDQFEFLLGKGLNKVTGLETALLDFLKHHCPENKELFTLVALHFRLYHELALMWENE   91 (319)
Q Consensus        12  ~VRLltGIgRy~EM~YiFdiL~en~qFE~Ll~k~~~~~~~~L~tAlLdylKr~~P~d~Ek~~mvAL~FsM~rEige~~e~~   91 (319)
                       +-|++.|= =+-++-..|....-|..|..+. .|+--|+--+.--..-. +..-+|..+|...+++
T Consensus      161 v~rl~~g~----VLLfLk~TL~~~~F~~lL~~~P--------~A~~~Yi~YL~~~~~~~------~L~~ll~~lgr~~eaA   223 (462)
T XP_011535368.1 161 VRRLRKGK----VLIFLKRTLSKEILFRELEVRQ--------VALRHLIHFLKEIGDQK-----LLLDLFRFLDRTEELA   223 (462)
T ss_pred               HHHHHHHH----HHHHHHHhCCHHHHHHHHHHhCH--------HHHHHHHHHHHHcCCHH-----HHHHHHHHCCChHHHH

Q ss_pred               HHHHHHHHHHhhhhHHHhhhccHhhHhhccHHHHHHHHHHHHHHHHHHHHHHHHhHHHHHHHHHHHHHHHHHHHHHHHhc
Q Apis_Spg11       92  AKDLIKTIISNATKEYNKLQSNVQHEIKFIKTEYIQKQLQLIITNYTHATEYYLQANKLNLASQCSDQVQLVALQLSLFN   171 (319)
Q Consensus        92  A~~~lk~I~sq~~~~~~~~Lq~~~~~~~~~~~~~~~~~~~kk~L~~~~t~~tdAaE~Y~kd~c~r~A~qc~~~a~Lv~lQl~lln   171 (319)
                       ...+-..+.....+. +++. |+++...|..+ ..+.......+|..+|+.+-.
T Consensus      224 ~l~~~~a~~~~~~~~~~--r~~~-----------------Lk~a~~~~~~~~----------~~~~~v~e~i~LLe~Q~~le~   274 (462)
T XP_011535368.1 224 LSHYREHLNIQDPDK--RKEF-----------------LKTCVGLPFSA----------EDSAHIQDHYTLLERQIIIEA   274 (462)
T ss_pred               HHHHHHHHHhcCChHH--HHHH-----------------HHHHhcCCCH----------hHHHHHHHHHHHHHHHHHHHHHH

Q ss_pred               ------------------cCCCCcceEEeecCC-HHHHHHHHHcCC-----chHHHHHHHHHHhcc-CCcHHHHHHHHHHH
Q Apis_Spg11      172 ------------------TTSYNQQVICILNLK-PEDIDKVLCHNL-----SFSQCFIIVHAYNH-HVDWANLIYNHCIL   226 (319)
Q Consensus       172 ------------------~~~~~~~~~~~~~~~~Inl~-~~~l~~~~~~~~~~~~~-----~f~Qa~iva~AY~~-~~dWae~iy~qvI~   226 (319)
                       . ...++|.+ .+-|..|+..|. .+..+.-++..|+. +--+.-.-.+-.+.
T Consensus      275 ~d~~~~~~~~~~~~~~f~~~p~------~~~~~vg~Sl~eTL~y~~~~h~~~~~~~~~~~~sp~~lkk~FkIsdKqf~w~~L~alAk   348 (462)
T XP_011535368.1 275 NDRHLESAGQTEIFRKHPR------KASILNMPLVTTLFYSCFYHYTEAEGTFSSPVNLKKTFKIPDKQYVLTALAARAK   348 (462)
T ss_pred               hHHHHHHcCCccchhHhCCC------cccCCCCCHHHHHHHHHHHhCcccCCCCcCCHHHHHHHcCCCHHHHHHHHHHHHHHH

Q ss_pred               hCCchHHHHHHHh-------hcCCCHHHHHHHHHHHHHhccCCcchHHHHHHHHHHHHHhhchHHHHHHHHhCcHHHH
Q Apis_Spg11      227 NGETKYLKDFIT-------VNKLTPSLVEDCVRRYRLEKSITHTMTNNMKILISELSDVECKYVLASQLGFKNIV   294 (319)
Q Consensus       227 ~g~f~YL~eF~~--------~~~l~~s~~edi~kky~~~kp~~~~~~~~~nmk~Li~~~~~dV~~~Y~LA~~lgF~~IV   294 (319)
                       .|+..-++.|.. ..+-+|=-++.++..+..++++ .+-+.+.+..|.|...++.||..+|-.++|
T Consensus      349 ~~~W~dLe~l~~~~K~~lg~~K~~spIg~~~~~v~~l~~~~~aP----~~~~l~kyl~~i~d~eeRl~la~k~~~~~~~a   419 (462)
T XP_011535368.1 349 LRAWNDVDALFTTKNWLGYTKKRAPIGFHRVVEILHKNNAP----VQILQEYVNLVEDVDTKLNLATKFKCHDVV   419 (462)
T ss_pred               ccCHHHHHHHHHcCCccccCCCCCCCCCCHHHHHHHHHHCCCC----HHHHHHHHHHhCCCHHHHHHHHHHHCCCHHHH
```

Template alignmentNCBI Fasta

10. NP_649877.1 vacuolar protein sorting 16A [Drosophila melanogaster]
Probability: 86.55%, E-value: 140, Score: 34.91, Aligned cols: 252, Identities: 15%, Similarity: 0.09,

```
Q ss_pred             HHHHHhchHHHHHHccC-CCccccCHHHHHHHHHHHHCCCChHHHHHHHHHHHHHHHHHHHhHHHHHHHHHHHHHHHHhhhhHH
Q Apis_Spg11  29   FQILKENDQFEFLLGKG-LNKVTGLETALLDFLKHHCPENKELFTLVALHFRLYHELALMWENEAKDLIKTIISNATKEY   107 (319)
Q Consensus  29   FdiL~en~qFE~Ll~k~-~~~~~~~~L~tAlLdylKr~~P~d~Ek~~mvAL~FsM~rEige~~e~~A~~~lk~I~sq~~~~~~   107 (319)
                     +..|.+-.+.+..|.+- ....+.+=-.++-++++..+ .++.+.+..-+=.........++..-.+.++.+-.+.-..
T Consensus    547 v~lLl~l~~~~~AL~~Ai~s~D~dlI~~vll~l~~~l~-~~~f~~~L~~~p~A~~~y~~ylr~~~~~~~L~~~~~~~~d~~-   624 (833)
T NP_649877.1 547 VPLLLKMRKFDRAVGSATQSGDTELITQVLLEMKMHMM-LSNLHMTIRDHPLALNIYKKIMRESNRAGLYGIYNTEDDQ-   624 (833)
T ss_pred             HHHHHhcccHHHHHHHHHHHcCCHHHHHHHHHHHHHHhhcC-HHHHHHHHHhCHHHHHHHHHHHHHHHcCHHHHHHHHHhccCcH-

Q ss_pred             HhhhccHhhHhhccHHHHHHHHHHHH------HHHHHHHHHHHHhhHHHHHHHHHHHHHHHHHHHHHHHHHhccCCCCcceEE
Q Apis_Spg11  108  NKLQSNVQHEIKFIKTEYIQKQLQLII------TNYTHATEYYLQANKLNLASQCSDQVQLVALQLSLFNTTSYNQQVIC   181 (319)
Q Consensus  108  ~~Lq~~~~~~~~~~~~~~~~~~~~~~~~kk~L~~~~------t~~tdAaE~Y~kd~c~r~A~qc~~~a~Lv~lQl~lln~~~~~~~~~~~~~~   181 (319)
                     .+-....++.+. .....|.+.|.....-..+..+.+..+|..+|..+-+... ..
T Consensus    625  ---------------~~~a~~~~~~~~~~~~~~~~~~~~~L~~~~~~~~~~~~~~~~~~~~~~~~~~~~~~e~~~Ll~~Q~~le~~~~~------~~   684 (833)
T NP_649877.1 625  ---------------KAIAEYHFQNAIETEGLESNLSMIGNAYAQGRCTLEAELCADTSRLIKLQKTLSTKYN-----VS   684 (833)
T ss_pred             ---------------HHHHHHHHHHHHHhcccHHHHHHHHHHHHHhcCCchHHHHHHHHHHHHHHHHHHHHHHHHhhhC-----CC

Q ss_pred             eecCC-HHHHHHHHHcCCchHHHHHHHHHHhcc-CCcHHHHHHHHHHHHhCCchHHHHHHHhcCCCHHHHHHHHHHHHhccCC
Q Apis_Spg11  182  ILNLK-PEDIDKVLCHNLSFSQCFIIVHAYNH-HVDWANLIYNHCILNGETKYLKDFITVNKLTPSLVEDCVRRYRLEKS   259 (319)
Q Consensus  182  ~Inl~-~~~l~~~~~~~~~~~~~~f~Qa~iva~AY~~-~~dWae~iy~qvI~~g~f~YL~eF~~~~~~~l~~s~~edi~kky~~~kp   259 (319)
                     +++++ .+-+..++..+ .+..|--++..++. +--|.-.-.+-.+..|++.-++.|.... -+|=-++.+++.+..++|
T Consensus    685  ~v~~Sl~~Tl~~l~~~~-~~~~a~~lkk~fkVsdK~~~~~kl~ala~~~~w~~L~~~~~~~k-kspIg~~~fv~~~~~~~~~~   762 (833)
T NP_649877.1 685  LNGLSIHETIQELLLIG-ELKEAERIRSEFKVPDRRFWWLRILTLSSQHKWEELEKLAKSK-KSPIGYDPFVEVCLKQDN   762 (833)
T ss_pred             cCCCCHHHHHHHHHHcC-CHHHHHHHHHHHHcCCChHHHHHHHHHHHHHHhcCCHHHHHHHHHhcC-CCCCCcHHHHHHHHHCCC

Q ss_pred             ccHHHHHHHHHHHHHHHHhhcHHHHHHHHHCcHHHHHHHHHhCcccchhhhhh
Q Apis_Spg11  260  ITHTMTNNMKILISELSDVECKYVLASQLGFKNIVEAMLNNPMIGAYLKDT   310 (319)
Q Consensus  260  ~~~~~~~~~nmk~Li~~~~~dV~~~Y~LA~~lgF~~IV~~LLnd~~~g~~LkDt   310 (319)
                     + +...+.|..|+|. -+++++.+.|-.+.+-...-.......-|...
T Consensus    763  ~-----~~a~~yI~~i~~~-~r~~l~~~~~~~~~~~~Aie~a~~~kD~~~L~~i   807 (833)
T NP_649877.1 763  V-----MEARKYIPRCRDN-RRVVWYMRANLFNEAIDSAFEQRDVHSLFEL   807 (833)
T ss_pred             H-----HHHHHHHHhCCcH-HHHHHHHHHCCCHHHHHHHHHHHcCCHHHHHHH
```

Template alignmentNCBI Fasta

11. NP_651731.1 vacuolar protein sorting 16B [Drosophila melanogaster]

Probability: 81.55%,  E-value: 180,  Score: 32.21,  Aligned cols: 242,  Identities: 10%,  Similarity: -0.048,

```
Q ss_pred             HHHHHHHhchHHHHHHccC-CCcccCHHHHHHHHHHHHHCCCC-----hHHHHHHHHHHHHHHHHHHHHhHHHHHHHHHHHHHH
Q Apis_Spg11   27  YIFQILKENDQFEFLLGKG-LNKVTGLETALLDFLKHHCPEN-----KELFTLVALHFRLYHELALMWENEAKDLIKTII   100 (319)
Q Consensus    27  YiFdiL~en~qFE~Ll~k~-~~~~~~L~tAlLdylKr~~P~d-----~Ek~~mvAL~FsM~rEige~~e~~A~~~lk~I~   100 (319)
                   |-++.+..-+.-+.||.+. ....+.+=.+++=+|++..+.. -.....|+=||.-|=.-...++ ...+++....
T Consensus   121 ~~L~~~rSl~eK~~lLd~Ai~s~DgdlI~~VllfLk~TL~~~~F~~lL~~~p~A~~~yi~yL~~~~~~~-~L~~~l~~lg   199 (447)
T NP_651731.1 121 APLEMFKSLHEKQQLLDTLMAQGGGHAVITVLLFLKRTLNTTQFHGILRERPKALEQYLSYLKESGDLA-GHIELLQLFG   199 (447)
T ss_pred             CCHHHhCCHHHHHHHHHHHHHcCCHHHHHHHHHHHHHhcCHHHHHHHHHhCHHHHHHHHHHHHHcCCHH-HHHHHHHHcC

Q ss_pred             HhhhhHHHhhhcchHhhHhhcchHHHHHHHH-----HHHHHHHHHHHHHHHh---hHHHHHHHHHHHHHHHHHHHHHHhhcc
Q Apis_Spg11  101 SNATKEYNKLQSNVQHEIKFIKTEYIQKQLQ-----LIITNYTHATEYYLQA---NKLNLASQCSDQVQLVALQLSLFNT   172 (319)
Q Consensus   101 sq~~~~~~~Lq~~~~~~~~~~~~~~~~kk~L~-----~~~t~~tdAaE~Y~kd---~c~r~A~qc~~~a~Lv~lQl~lln~   172 (319)
                 +..-.. -..++..++ .-.+....|.+.|.+. ..--.++.+....+|..+|..+-..
T Consensus   200 r~~eaA----------------~~~~~~a~~~~~~~~~r~~~L~~~~~~~~~~~~~~~~~~~~~~~v~~~i~LLe~Q~~l~~~   263 (447)
T NP_651731.1 200 RHQEAA----------------LKQFQAALASGDTSTRKKHLQLLVDAYATAGVGVIPLYEQVFHAALKMQQLMEKDSAL   263 (447)
T ss_pred             CHHHHH----------------HHHHHHHHHcCChHHHHHHHHHHHHHHHHHhcCCCChhHHHHHHHHHHHHHHHHHHHHHHhh

Q ss_pred             CCCCcceEEeecCC-HHHHHHHHHcC---C-----chHHHHHHHHHhcc-CCcHHHHHHHHHHHhCCchHHHHHHhhcCC
Q Apis_Spg11  173 TSYNQQVICILNLK-PEDIDKVLCHN---L-----SFSQCFIIVHAYNH-HVDWANLIYNHCILNGETKYLKDFITVNKL   242 (319)
Q Consensus   173 ~~~~~~~~~~~Inl~-~~~l~~~~~~~~---~-------~f~Qa~iva~AY~~-~~dWae~iy~qvI~~g~f~YL~eF~~~~~l   242 (319)
                 . ...++|.+ .+-|..|+..| . ++..+.-++..|+. +--+.-.-.+-....|+..-++.|......
T Consensus   264 ~-----~~~~~~~Sl~~TL~y~~~~h~~~~~~~~~~~~~~sp~~lkk~FkIseKqf~w~~L~ala~~~~W~~le~l~~~K~~   338 (447)
T NP_651731.1 264 G-----KLLRDQPTPVEVLYACCQVNSNWKEQDMLKPVSPQRFAADQQISPAQYEWTALNERAQAQAYADLECIFERVPS   338 (447)
T ss_pred             c-----cccCCCCCHHHHHHHHHHHcCCCcccccCCCCCHHHHHHHcCCCHHHHHHHHHHHHHHHHHhcchHHHHHHHHhcCCC

Q ss_pred             ----------CHHHHHHHHHHHHhccCCcchHHHHHHHHHHHHHHHhhchHHHHHHHHHhCcHHHH
Q Apis_Spg11  243 ----------TPSLVEDCVRRYRLEKSITHTMTNNMKILISELSDVECKYVLASQLGFKNIV   294 (319)
Q Consensus   243 ----------~~s~~edi~kky~~~kp~~~~~~~~~nmk~Li~~~~dV~~~Y~LA~~lgF~~IV   294 (319)
                 .|=-++-++.....++++ .+-+.+.+..|.|...++.||...|-.++|
T Consensus   339 ~lg~~kk~~~~pig~~~~~v~~l~~~~ap----~~~l~kyi~~i~d~e~Rl~la~k~~~~~~a   396 (447)
T NP_651731.1 339 WHPLKTKQFHISFDLTLAVLRLYELQAP----STVLQLFLSKMGNSVEKLALAQRVKCIKAV   396 (447)
T ss_pred             cCCCCcccccCCCCHHHHHHHHHHHCCCC----HHHHHHHHHcCCCHHHHHHHHHHHHccCHHHH
```
